# Pan-cancer single cell RNA-seq uncovers recurring programs of cellular heterogeneity

**DOI:** 10.1101/807552

**Authors:** Gabriela S. Kinker, Alissa C. Greenwald, Rotem Tal, Zhanna Orlova, Michael S. Cuoco, James M. McFarland, Allison Warren, Christopher Rodman, Jennifer A. Roth, Samantha A. Bender, Bhavna Kumar, James W. Rocco, Pedro ACM Fernandes, Christopher C. Mader, Hadas Keren-Shaul, Alexander Plotnikov, Haim Barr, Aviad Tsherniak, Orit Rozenblatt-Rosen, Valery Krizhanovsky, Sidharth V. Puram, Aviv Regev, Itay Tirosh

## Abstract

Cultured cell lines are the workhorse of cancer research, but it is unclear to what extent they recapitulate the cellular heterogeneity observed among malignant cells in tumors, given the absence of a native tumor microenvironment. Here, we used multiplexed single cell RNA-seq to profile ~200 cancer cell lines. We uncovered expression programs that are recurrently heterogeneous within many cancer cell lines and are largely independent of observed genetic diversity. These programs of heterogeneity are associated with diverse biological processes, including cell cycle, senescence, stress and interferon responses, epithelial-to-mesenchymal transition, and protein maturation and degradation. Notably, some of these recurrent programs recapitulate those seen in human tumors, suggesting a prominent role of intrinsic plasticity in generating intra-tumoral heterogeneity. Moreover, the data allowed us to prioritize specific cell lines as model systems of cellular plasticity. We used two such models to demonstrate the dynamics, regulation and drug sensitivities associated with a cancer senescence program also observed in human tumors. Our work describes the landscape of cellular heterogeneity in diverse cancer cell lines, and identifies recurrent patterns of expression heterogeneity that are shared between tumors and specific cell lines and can thus be further explored in follow up studies.

## Introduction

Cellular plasticity and heterogeneity are fundamental features of human tumors driven by both genetic and epigenetic mechanisms (Chaffer et al., 2016; McGranahan and Swanton, 2015). Malignant cells within a single tumor display diverse patterns of gene expression, which underlie differences in morphology, metabolism, proliferation, invasion and immunogenicity. The existence of cells with multiple phenotypes plays a major role in disease progression and treatment failure, as subpopulations of cells may drive tumor recurrence and metastasis. Thus, it is critical for cancer research to establish frameworks to characterize cellular diversity within tumors, as well as the underlying mechanisms that generate such diversity.

Single-cell RNA sequencing (scRNA-seq) has emerged as a valuable tool to study cell-to-cell heterogeneity within tumors (Chung et al., 2017; Filbin et al., 2018; Kim et al., 2016; Lambrechts et al., 2018; Li et al., 2017; Patel et al., 2014; Puram et al., 2017; Tirosh et al., 2016a; Tirosh et al., 2016b; Venteicher et al., 2017). Several studies identified functionally significant patterns of intra-tumoral heterogeneity (ITH) within malignant cells, yet their origin and mechanisms were difficult to resolve from observations in patients.

In principle, genetic diversity, epigenetic cell-intrinsic plasticity, and interactions with the spatially-variable tumor microenvironment all contribute to the heterogeneity observed across malignant cells. However, since previous studies suggest that major patterns of expression heterogeneity in tumors are linked to their cell-of-origin and recapitulated in cell lines, we hypothesize that they may reflect intrinsic cellular plasticity that exists even in the absence of genetic diversity and a native microenvironment. For example, we previously reported an epithelial-to-mesenchymal transition (EMT)-like program associated with metastasis in head and neck squamous cell carcinoma (HNSCC) that was partly preserved in one of a number of tested cell lines (Puram et al., 2017). Similarly, drug resistance melanoma programs identified in tumors were recapitulated and studied in melanoma cell lines (Jerby-Arnon et al., 2018; Shaffer et al., 2017; Tirosh et al., 2016a).

Human cell lines are a mainstay of cancer research and drug discovery, yet our current knowledge of their ability to recapitulate the expression diversity observed in patient samples is limited. Only a few cancer cell lines have been comprehensively profiled by scRNA-seq so far (Ben-David et al., 2018; Jerby-Arnon et al., 2018; Kim et al., 2015; Sharma et al., 2018). Thus, models are often chosen based on their mutational status, historical popularity, and ease of culturing.

Here, we apply multiplexed scRNA-seq to profile 198 cell lines from 22 tumor types in the Cancer Cell Line Encyclopedia (CCLE) collection (Barretina et al., 2012; Ghandi et al., 2019). Analysis of expression heterogeneity within cell lines revealed patterns of variability that recurred across different cell lines, and spanned diverse biological functions. Strikingly, many of these variable programs observed *in vitro* matched those previously characterized in patient tumors. We used these results to select model cell lines and utilize them for follow up studies, demonstrating the dynamics, regulation, and drug sensitivities associated with a recurrent expression program linked to cellular senescence. Likewise, this dataset provides a resource for the rational selection of cancer cell lines as models for exploring the determinants and consequences of ITH.

## Results

### Pan-cancer scRNA-seq of human cell lines

To effectively profile expression heterogeneity within diverse cancer cell lines, we developed and applied a multiplexing strategy where cells from different cell lines are grown and profiled in pools and then computationally assigned to the corresponding cell line (Fig. 1A). We utilized existing pools that were previously generated from the CCLE collection (Barretina et al., 2012; Yu et al., 2016). Each pool consisted of 24-27 cell lines from diverse lineages but with comparable proliferation rates, and was profiled by massively parallel scRNA-seq, for an average of 280 cells per cell line (**Methods**). We profiled eight CCLE pools, along with one smaller custom pool that included HNSCC cell lines.

**Figure 1.**
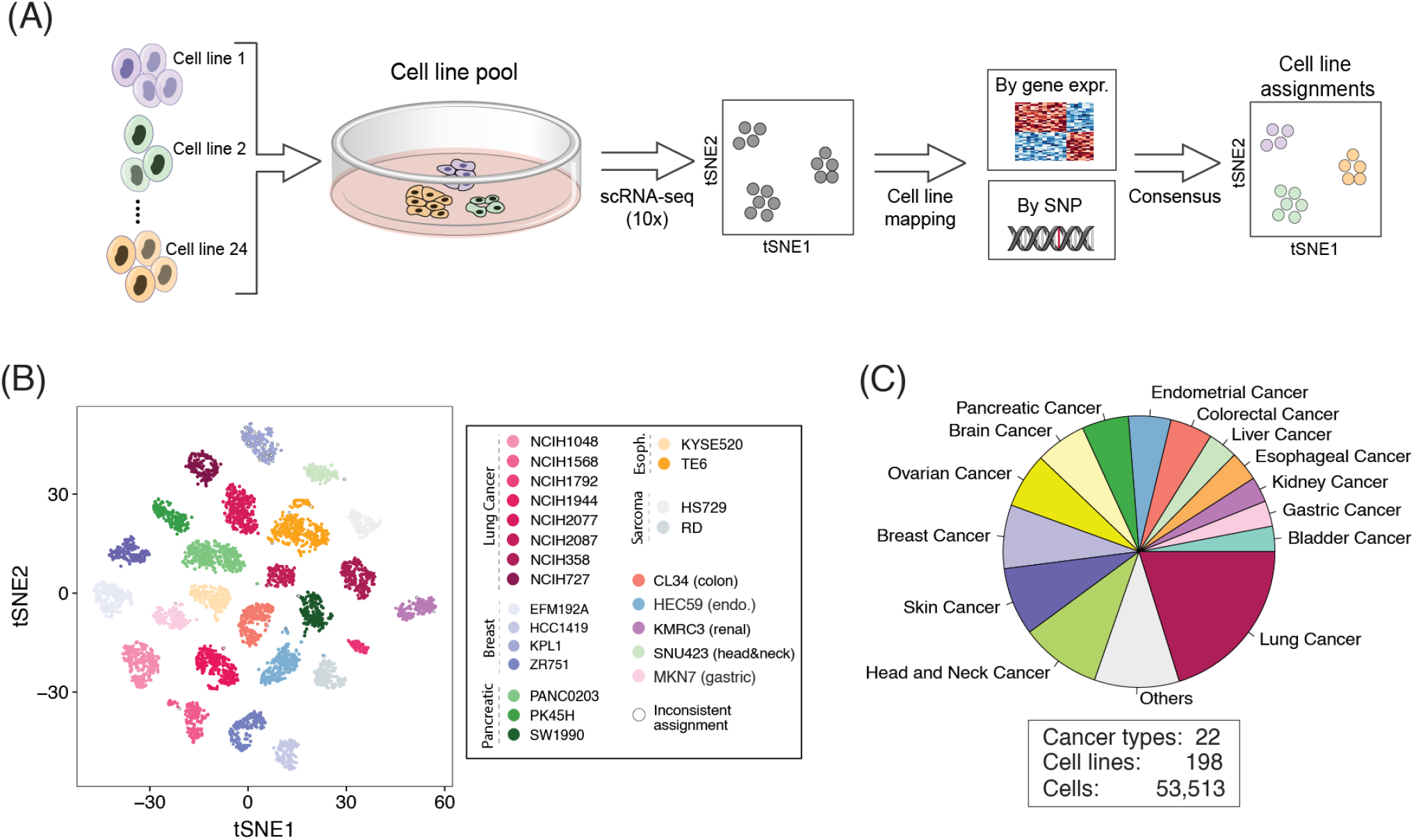
Characterizing intra-cell line expression heterogeneity by multiplexed scRNA-seq. (**A**) Workflow of the multiplexing strategy used to profile multiple cell lines simultaneously. Cell lines were pooled and profiled by droplet-based scRNA-seq. We used reference CCLE data to assign cells to the most similar cell line based on their overall gene expression and SNP pattern. (**B**) t-SNE plot of a representative pool demonstrating the robustness of cells’ assignments to cell lines. Cells with inconsistent assignments (by gene expression and SNPs) are denoted and these were excluded from further analyses. (**C**) Distribution of cancer types profiled.

We assigned profiled cells to cell lines based on consensus between two complementary approaches, using genetic (SNP) and expression profiles (Fig. 1A). First, cells were clustered by their global expression profile, and each cluster was mapped to the cell line with the most similar bulk RNA-seq profile (Ghandi et al., 2019). Second, by detection of SNPs in the scRNA-seq reads, we assigned cells to the cell line with highest similarity by SNP profiles derived from bulk RNA-seq (Ghandi et al., 2019; Kang et al., 2018). Cell line assignments based on gene expression and SNPs were consistent for 98% of the cells, which were retained for further analysis (*e.g.* Fig. 1B). The few inconsistent assignments were observed primarily in cells with low data quality, resulting in low SNP coverage, which were therefore excluded. Cell lines with less than 50 assigned cells were also excluded from further analyses, as were low-quality cells and suspected doublets.

Overall, following assignment and quality control filters, we studied the expression profiles of 53,513 cells, from 198 cell lines (56-1,990 cells per cell line; **fig. S1A**), reflecting 22 cancer types (Fig. 1C). We detected an average of 19,264 UMIs and 3,802 genes per cell, underscoring the high quality of our dataset (**fig. S1A**).

### Discrete and continuous patterns of expression heterogeneity within cell lines

We aimed to characterize the variation in gene expression across cells within individual cell lines, distinguishing between discrete patterns, reflecting distinct subpopulations of cells, and continuous patterns, reflecting spectra of cellular states (Fig. 2A).

**Figure 2.**
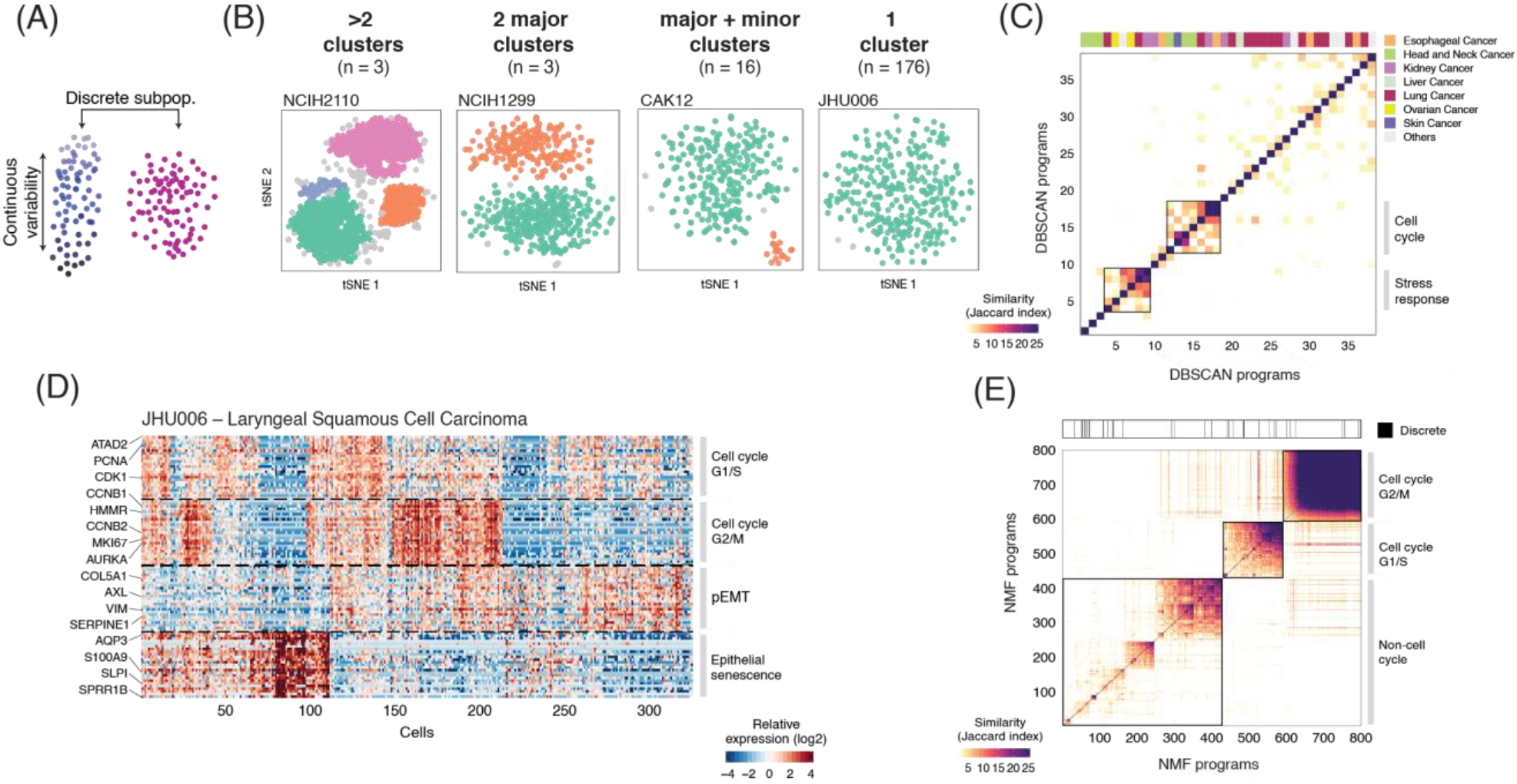
Discrete and continuous patterns of heterogeneity within cell lines. (**A**) Illustration of the two types of expression variability investigated. (**B**) t-SNE plots show exemplary cell lines for the four classes defined by the presence and number of discrete subpopulations identified using DBSCAN. The description of each class and number of cell lines is indicated above the t-SNE plots. (**C**) Heatmap depicts pairwise similarities between gene expression programs defined for each of the cell clusters derived from the 22 cell lines identified as having one or more discrete subpopulations. Hierarchical clustering identifies only two groups of similar programs (metaprograms). Top panel shows assignment to cancer types. (**D**) Continuous programs of heterogeneity identified using NMF in a representative cell line that lacks discrete subpopulations (JHU006; see **B**). Heatmap shows relative expression of genes from four programs, across all cells ordered by hierarchical clustering. NMF programs are annotated (right) and selected genes are indicated (left). (**E**) Pairwise similarities between NMF programs identified across all the cell lines analyzed and ordered by hierarchical clustering. Programs with limited similarity to all other programs as well as those associated with technical confounders were excluded. Top panel indicates the 4% of NMF programs that were consistent with discrete subpopulations identified by DBSCAN (*P*<0.001, Fisher’s exact test).

To identify discrete subpopulations we used dimensionality reduction with t-Distributed Stochastic Neighbor Embedding (t-SNE) followed by density-based clustering (DBSCAN; **fig. S1B; Methods**). Discrete clusters of cells within a cell line were found only for a minority (11%) of the cell lines: three cell lines had three or more clusters, three had two clusters of comparable sizes, and 16 had one major and one minor cluster (Fig. 2B and **S1C**). For each such cluster, we identified the top 50 upregulated genes compared to all other cells from the same cell line. These expression programs showed limited similarities to one another, both within cell lines of the same cancer type and across different cancer types, indicating that discrete subpopulations are typically unique and cell line-specific (Fig. 2C). The main exceptions were seven subpopulations commonly upregulating cell cycle-related genes, and six subpopulations commonly upregulating stress response genes. Similar results were obtained using DBSCAN with different parameters (**fig. S1D-E**).

To also identify continuous variability of cellular states, we applied non-negative matrix factorization (NMF) to each cancer cell line (Puram et al., 2017). We repeated the NMF analysis with distinct parameters, to identify robust expression programs (*i.e.* consistently observed as variable using different parameters), each defined by the top 50 genes based on NMF scores (*e.g.*, Fig. 2D; **Methods**). This procedure captures both continuous and discrete programs. Overall, we detected 1,445 robust expression programs across all cell lines, with 4-9 such programs in individual cell lines (**fig. S1F**). To identify common expression programs varying within multiple cell lines, we first excluded those with limited similarity to all other programs as well as those associated with the technical confounder of variable cell quality (**fig. S1G**), retaining 800 programs (0-8 per cell line, **fig. S1F**). Of these programs, only 4.75% corresponded to the discrete subpopulations above (Fig. 2E).

Hierarchical clustering of the NMF programs based on their shared genes emphasized multiple recurrent heterogeneous programs (RHPs) of gene expression, which are present in multiple cell lines. The two most prominent RHPs reflected the cell cycle, and 10 additional RHPs were associated with other cellular processes (Fig. 2E). The cell cycle RHPs corresponded to the G1/S and the G2/M phases (Fig. 2E), as was also observed in clinical tumor samples (**Fig. S2A**). G2/M programs were similar across cell lines, as well as between cell lines and tumors, thus defining a generic mitotic program (**fig. S2B**). In contrast, G1/S programs differed more both across cell lines and between cell lines and tumors (**fig. S2B**), indicating that expression programs associated with genome replication are more context-dependent. A central difference in G1/S programs involved the MCM complex genes (MCM2-7) and the linker histone H1 family genes (HIST1H1B-E), which were robustly upregulated only in tumors or cell lines, respectively (**fig. S2B,D**). This may reflect an *in vitro* adaptation to rapid growth and loss of the G1 checkpoint in cell lines. Consistent with this possibility, while tumors have a high percentage of apparent G0 cells (*i.e.*, lacking both G1/S and G2/M expression programs), such cells are much less prevalent in cell lines (**fig. S2E**).

### RHPs reflect distinct biological processes and mirror *in vivo* states

The ten additional (non-cell cycle) RHPs reflected diverse biological processes, and are described further below. Importantly, several analyses indicated that the pooling approach had limited impact on defining heterogeneity within individual cell lines, and that RHPs were robustly identified across pools. First, the similarity between NMF programs was largely independent of the pools in which they were detected (**fig. S3A**). Second, RHPs were each detected across at least four different pools (**fig. S3B**). Third, a single cell line had highly similar NMF programs when profiled in two distinct pools (**fig. S3C**).

We next characterized these 10 RHPs by functional enrichment of their signature genes, the lineages and mutations of cell lines in which they are observed, as well as their similarity with programs that vary across cells in patient tumor samples (Fig. 3A-C). Overall, 7 out of 10 RHPs resemble the heterogeneity observed in human tumor samples, including 5 RHPs with particularly high similarity (Fig. 3B, **S4A,B**), demonstrating the *in vivo* relevance of cell line RHPs.

**Figure 3.**
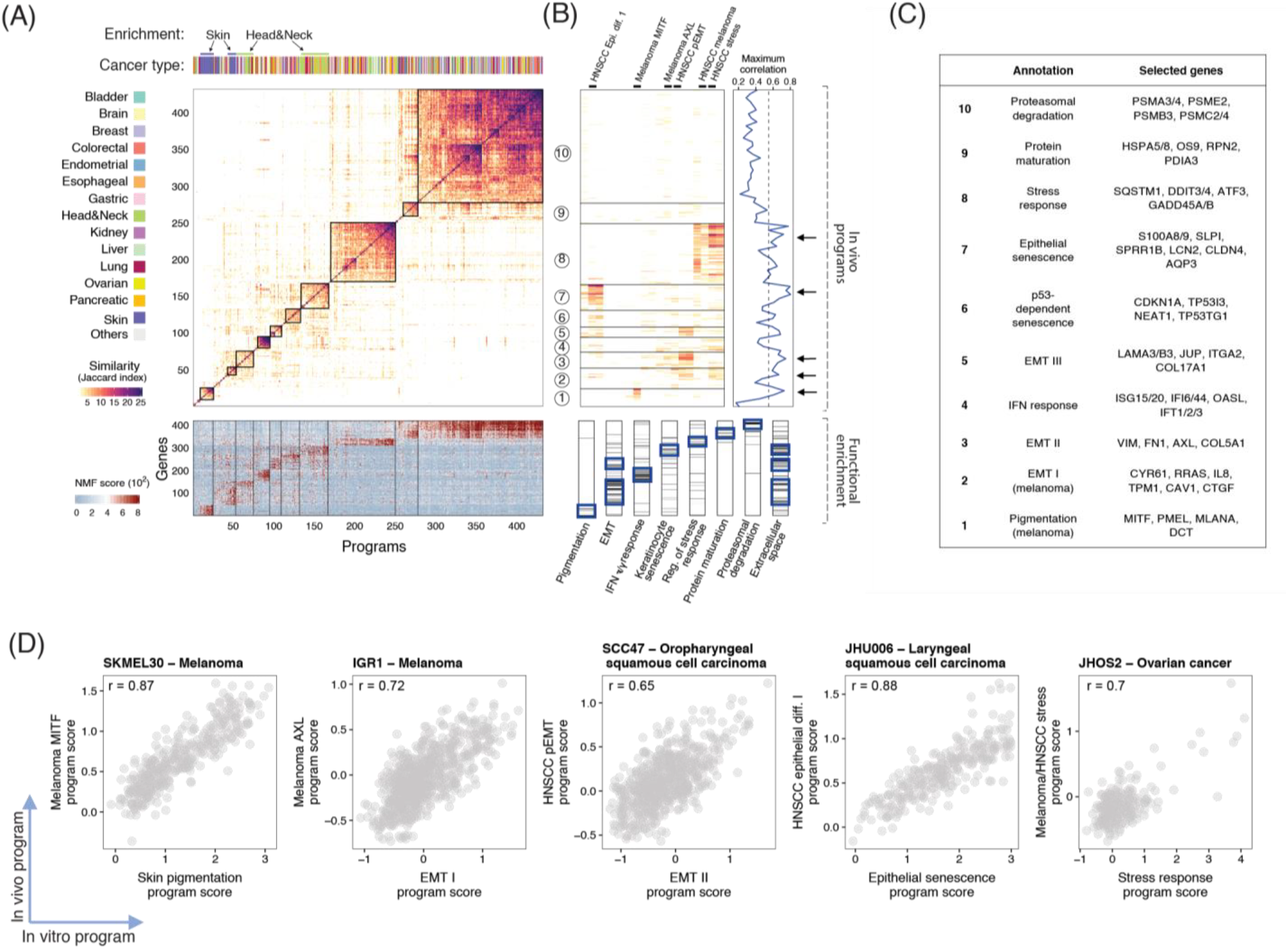
Functional annotation of RHPs. (**A)** The main heatmap depicts pairwise similarities between all NMF programs (except for those linked to the cell cycle, see Fig. 2E), ordered by hierarchical clustering. Ten clusters (RHPs) are indicated by squares and by numbers on the left. Top panel shows assignment to cancer types, highlighting significant enrichment (*P*<0.05, hypergeometric test) of melanoma and HNSCC cell lines. Bottom panel shows NMF scores of signature genes of each RHP. (**B**) **Top**: heatmap depicts similarities between NMF programs identified in cell lines (*in vitro*) and those observed in tumor samples (*in vivo*). Line plot on the right shows the maximum correlation of each *in vitro* program, emphasizing 5 RHPs (see **fig. S4A,B**) that closely recapitulate *in vivo* programs. Dashed line shows the maximum correlation obtained using 100 permutations. Bottom panel shows enrichment (*P*< 0.05, hypergeometric test) of RHP genes with eight annotated gene-sets. (**C)** Annotation and selected top genes for each of the 10 RHPs. (**D)** Single-cell scores of *in vitro* RHPs (X-axis) and the corresponding *in vivo* programs (Y-axis), in selected model cell lines.

One of the RHPs (#8) reflected a stress response, including DNA damage-induced and immediate early genes (*e.g.* DDIT3-4 and ATF3). This RHP resembles programs of heterogeneity previously observed in melanoma and HNSCC tumors (Fig. 3B,D) (Puram et al., 2017; Tirosh et al., 2016a), and may reflect the response to various cellular insults. Another RHP (#4) contained interferon (IFN) response genes (*e.g.*, IFT1-3 and ISG15,20). Recent studies revealed that IFN response may be triggered by genomic instability through the cGAS-STING pathway (Chen et al., 2016). Accordingly, the IFN-response program was depleted in cell lines with mutations in MRE11A (**fig. S5A**), which recognizes cytosolic dsDNA and activates STING (Kondo et al., 2013). Two other RHPs (#9 and #10) consisted of genes related to protein folding and maturation (e.g. HSPA1A, RPN2) and to proteasomal degradation (e.g. PSMA3-4), respectively. These RHPs, as well as the IFN response RHP #4, did not resemble the *in vivo* programs of heterogeneity observed previously among tumor cells. However, it is possible that such programs exist *in vivo* and have not been detected yet due to the limited scRNA-seq data in tumors.

### RHPs recapitulate *in vivo* EMT programs, and are associated with specific cancer types and NOTCH mutations

Three distinct RHPs were related to EMT: two shared across cancer types, and one unique to melanoma cell lines. The melanoma-specific EMT (RHP #2; EMT-I) was negatively correlated with another melanoma-specific RHP (#1) that was enriched with skin pigmentation genes (*e.g.*, MITF and PMEL). Both of these melanoma-specific RHPs, and their negative correlation, recapitulated the patterns of variability previously observed in melanoma tumors (Fig. 3B,D, **S4B,C**), in which they were linked to drug resistance (Shaffer et al., 2017; Tirosh et al., 2016a). Notably, as observed in patient samples, many of the melanoma cell lines (50%) harbored cells in both of these alternate cellular states, yet our data highlight certain melanoma cell lines as faithful model systems for these *in vivo*-related RHPs (**fig. S5C, S4C**).

Two other RHPs, EMT-II (#3) and EMT-III (#5), also reflected EMT-like processes in distinct cell lines. EMT-II was mainly observed in HNSCC cell lines (Fig. 3A), although across 7 distinct pools (**fig. S3B**). It included vimentin (VIM), fibronectin (FN1), the AXL receptor tyrosine kinase, and other genes, closely mirroring the partial EMT state we previously observed in HNSCC tumors (Fig. 3B,D, **S4B,C**), where it was linked to metastasis (Puram et al., 2017). Cell lines harboring EMT-II were depleted of *NOTCH4* mutations (**fig. S3B**) and were sensitive to inhibitors of NOTCH signaling (**fig. S5D**), suggesting a potential role of the NOTCH pathway in enabling EMT-II variability. This is similar to the association we found in glioblastoma between specific mutations and patterns of intra-tumoral heterogeneity (Neftel et al., 2019). In contrast, EMT-III was enriched among non-cycling cells (**fig. S5B**) and contained genes involved in cell junction organization such as laminin A3, B3 and C3, and plakoglobin (JUP). Interestingly, JUP was shown to promote collective migration of circulating tumor cells with increased metastatic potential (Aceto et al., 2014). The identification of three distinct EMT programs, two of which are enriched in specific cancer types, highlights EMT as a common, yet context-specific, pattern of cellular heterogeneity, which may have important implications for metastasis and drug responses.

### RHPs recapitulate classical and epithelial senescence programs

RHPs #6 and #7 were preferentially observed in G0 cells (**fig. S5B**) and seem to reflect different expression programs of cellular senescence. RHP #6 was enriched in p53-wild type cell lines and in those sensitive to the pharmacological activation of p53 by the MDM2 inhibitor Nutlin-3a (**fig. S5A,D**). Moreover, it included the senescence mediator p21 (CDKN1A) and other p53-target genes. Thus, we annotated it as “classical” p53-associated senescence. In contrast, RHP #7 was enriched in HNSCC cell lines, and was highly similar to the senescence program of keratinocytes (Hernandez-Segura et al., 2017) (Fig. 3A,B). RHP #7 also contained many secreted factors, such as S100A8/9, SAA1/2, LCN2, SLPI, which are involved in inflammatory responses and are reminiscent of the Senescence-Associated Secretory Phenotype (SASP).

To further examine the possibility that RHP #7 reflects a senescence program despite the lack of classical markers (*e.g.*, p16 and p21), we profiled primary lung bronchial cells by bulk RNA-seq after induction of senescence by etoposide. The etoposide-treated cells stained for the senescence marker SA-β-GAL (**fig. S5F**) and, compared to control, upregulated the expression of genes of both senescence-associated RHPs (#6 and #7) and downregulated the expression of cell cycle genes (**fig. S5E**).

Hence, RHP #7 resembles the senescence response of both keratinocytes and lung bronchial cells, while it differs from published senescence signatures of fibroblasts and melanocytes (Hernandez-Segura et al., 2017), underscoring the context- and cell type-specificity of senescence expression programs. We therefore denote it as an epithelial senescence (EpiSen) program. Notably, the EpiSen RHP recapitulates a program we previously observed in HNSCC tumors (fig. 3B,D, **S4B,C**), which was also negatively associated with cell cycle and spatially restricted to the hypoxic tumor core (Puram et al., 2017). We conclude that programs of cellular senescence are observed in subpopulations of cells in tumors and in cell lines and are associated with distinct expression profiles depending on genetics (*e.g.*, p53 status), lineage (*e.g.*, HNSCC), and possibly other features.

### Assessing the role of genetic heterogeneity through inference of chromosomal aberrations

Expression heterogeneity within a single cell line could be driven by either genetic or nongenetic mechanisms. To search for the contribution of genetic heterogeneity, we first identified genetic subclones by inferring large-scale copy number aberrations (CNAs) from the gene expression patterns in windows of 100 genes around each locus (Tirosh et al., 2016b). The inferred CNA profiles were consistent with hallmark genomic alterations such as the gain of chromosome 7 and loss of chromosome 10 in most glioblastoma cell lines (**fig. S6**). Importantly, CNA analysis allowed the robust identification of genetic subclones in 58 cell lines, based on the gain or loss of chromosomes (or chromosome arms) in a subset of cells (**Methods**).

Next, we compared the assignment of cells to genetic subclones with their patterns of expression heterogeneity. Twelve of the 22 cell lines (54%) with discrete expression-based clusters have genetic subclones, and 66% of their expression-based clusters were significantly correlated with genetic subclones (*P*<0.001, Fisher’s exact test; Fig. 4A). Thus, 39% of all discrete clusters were significantly associated with identified genetic subclones (Fig. 4B). Consistencies between genetic-based and expression-based classifications were much more limited for the continuous patterns of expression variability identified by NMF. Genetic subclones were observed only in 29% of the cell lines with continuous programs. Among these cell lines, only 31% of continuous programs were differentially expressed between genetic subclones (*P*<0.001, t-test). Taken together, only 8% of all continuous programs (compared with 39% for discrete clusters) were significantly associated with identified genetic subclones (Fig. 4B). In summary, genetic diversity, as evaluated by CNA-based subclones, partially contributes to expression heterogeneity in cell lines, and this effect is particularly weak for the continuous programs, underscoring the potential role of nongenetic regulation.

**Figure 4.**
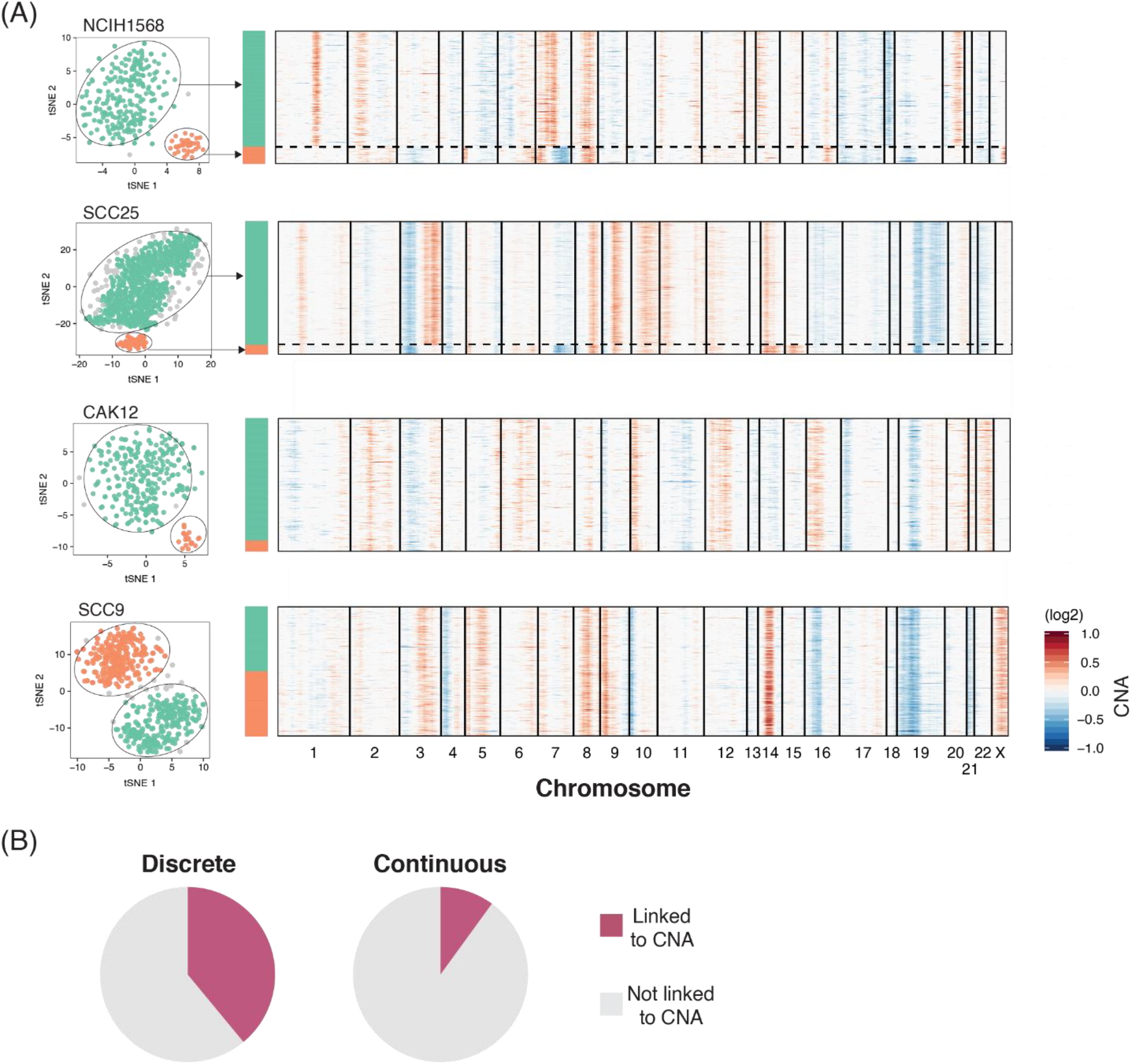
Genetic subclones partially explain expression heterogeneity. (**A)** Representative cell lines showing the association (top two cases) or lack thereof (bottom two cases) between discrete subpopulations and CNA-based subclones. t-SNE plots on the left show discrete subpopulations identified using DBSCAN (as in Fig. 2B and **S2B**). Heatmaps on the right depict inferred CNAs ordered according to the expression-based clusters. (**B)** Percentage of discrete (left) and continuous (right) heterogeneity programs that are associated with genetic subclones. For discrete programs, associations were assessed by comparing the assignment of cells to CNA subclones and to expression-based subpopulations (Fisher’s exact test *P*<0.001); for continuous programs, we compared NMF cell scores between different clones (*P*<0.001, t-test).

### Plasticity of heterogeneous cellular states in model cell lines

To assess the role of non-genetic mechanisms in regulating RHPs, we used two cell lines that successfully model both EMT-II (RHP #3) and EpiSen (RHP #7): JHU006, an HPV-laryngeal HNSCC, and SCC47, an HPV+ oropharyngeal HNSCC (Fig. 3D, **S4C**). Notably, EpiSen-high and EpiSen-low cells could be prospectively isolated, but returned to their pre-sorted heterogenous distribution with time. Specifically, we isolated by FACS EpiSen-high (AXL^−^/CLDN4^+^) vs. EpiSen-low (AXL^+^/CLDN4^−^) subpopulations, with ~12-fold difference in the expression of the EpiSen program (Fig. 5A-B). These sorted subpopulations began to shift by one week in culture and returned to the pre-sorting distribution of cellular states by day 14 (Fig. 5C; **fig. S7A**). The EpiSen-high subpopulation was enriched for G0/G1 phases, consistent with low proliferation (Fig. 5D; **fig. S7B**). Nevertheless, it still contained cells in the S and G2/M phases, and did not stain for the classical senescence marker SA-β-gal (data not shown). These results suggest that the EpiSen program is dynamically regulated by non-genetic processes, and that it does not represent a permanent exit from cell cycle, consistent with the observation that in cancer cells senescence is often an incomplete and reversible state (Lee and Schmitt, 2019).

**Figure 5.**
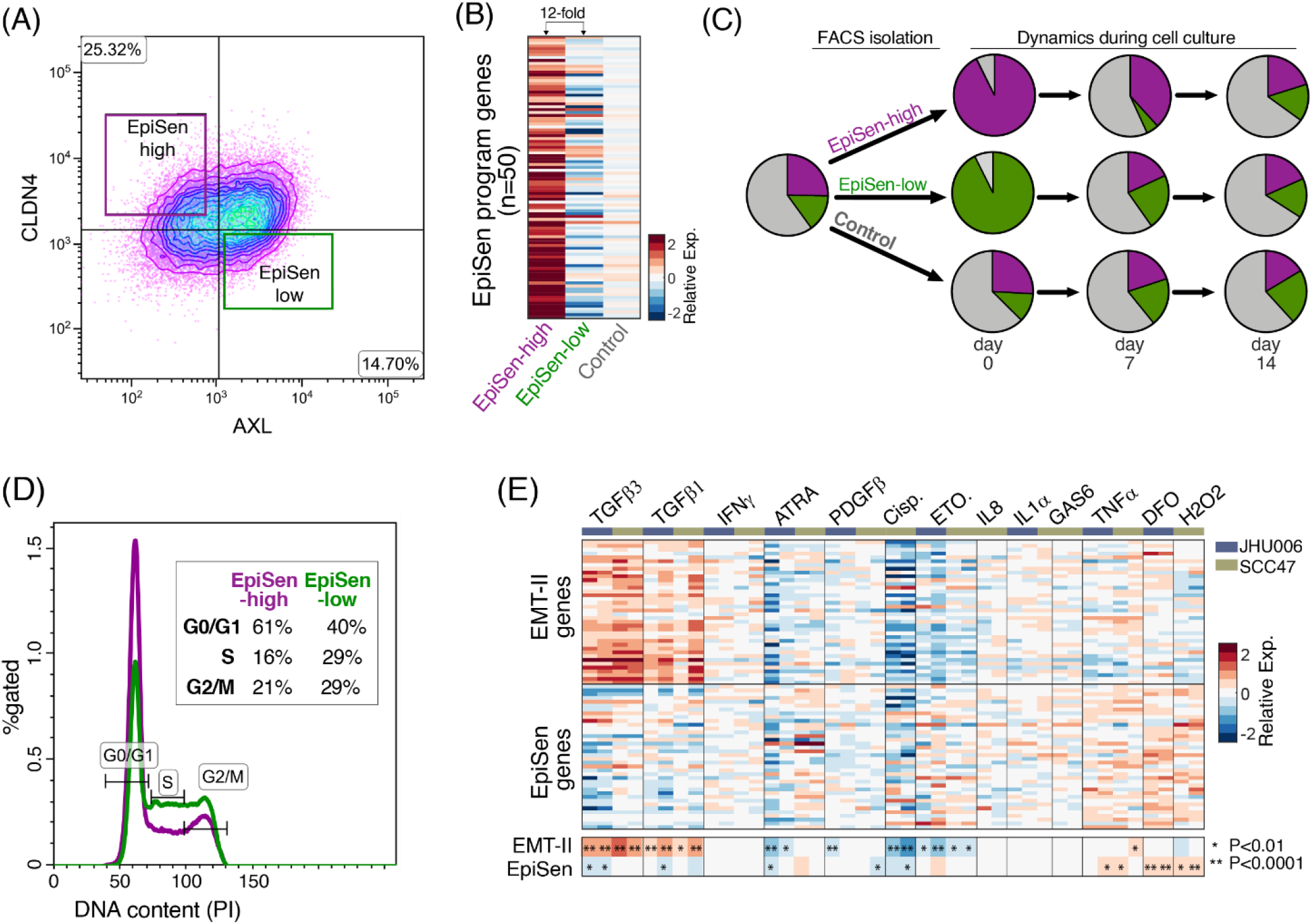
Interrogating the EpiSen RHP in model cell lines. **(A)** Isolation of the EpiSen-high (AXL^−^ CLDN4^+^) and EpiSen-low (AXL^+^CLDN4^−^) populations by FACS in JHU006. **(B)** Heatmap shows relative expression of the EpiSen program genes in three sorted subpopulations: two as shown in (A) and a third control population. **(C)** Pie charts depict relative proportions of the EpiSen-high and EpiSen-low subpopulations, for an unsorted sample (left, initial distribution), and for sorted subpopulations that were analyzed immediately after sorting (day 0) and at two additional time points (at days 7 and 14 in culture). **(D)** FACS analysis of cell cycle by the DNA binding dye propidium iodide (PI) on sorted EpiSen-high and EpiSen-low cells in JHU006. **(E)** Main heatmap depicts relative expression of EpiSen program genes and EMT-II program genes following multiple perturbations in SCC47 and JHU006. Smaller heatmap at the bottom shows the average values for the EMT-II genes and EpiSen genes, and asterisks denote significant up or down-regulation (by t-test).

Next, we examined the induction of these programs by tumor microenvironment soluble factors and perturbations (Fig. 5E). As expected, TGF-β1 and TGF-β3 upregulated the expression of the EMT-II genes, although the complete program induced had subtle differences from the native EMT-II program that was observed without perturbations and was associated with increased migration in a wound healing assay (**fig. S7C-D**). Interestingly, TGF-β treatments also downregulated the expression of EpiSen genes, underscoring the potential interplay between these two programs. A negative association between EMT and EpiSen was further supported by the single cell profiles of JHU006 and SCC47 cells (**fig. S4C**) and by our prior findings in HNSCC clinical samples, in which EMT-high cells were enriched at the invasive edge, while senescent cells were enriched at the core of tumors (Puram et al., 2017). Tumor cores are often associated with increased hypoxia, suggesting a potential mechanism for the spatial enrichment of senescent cells. In accordance with this possibility, the hypoxia mimetic desferrioxamine (DFO) induced the expression of the EpiSen program. A similar effect was observed upon hydrogen peroxide treatment, consistent with oxidative stress and the resultant DNA damage response as a potent inducer of senescence (te Poele et al., 2002) (Fig. 5E).

### Co-existing subpopulations differ in drug sensitivity

An important implication of cellular diversity in cancer is the possibility that distinct subpopulations of cells respond differently to treatments and thereby facilitate treatment failure and recurrence. Thus, we compared the sensitivities of EpiSen-high and EpiSen-low subpopulations sorted from each of the two model cell lines selected (Fig. 6A).

**Figure 6.**
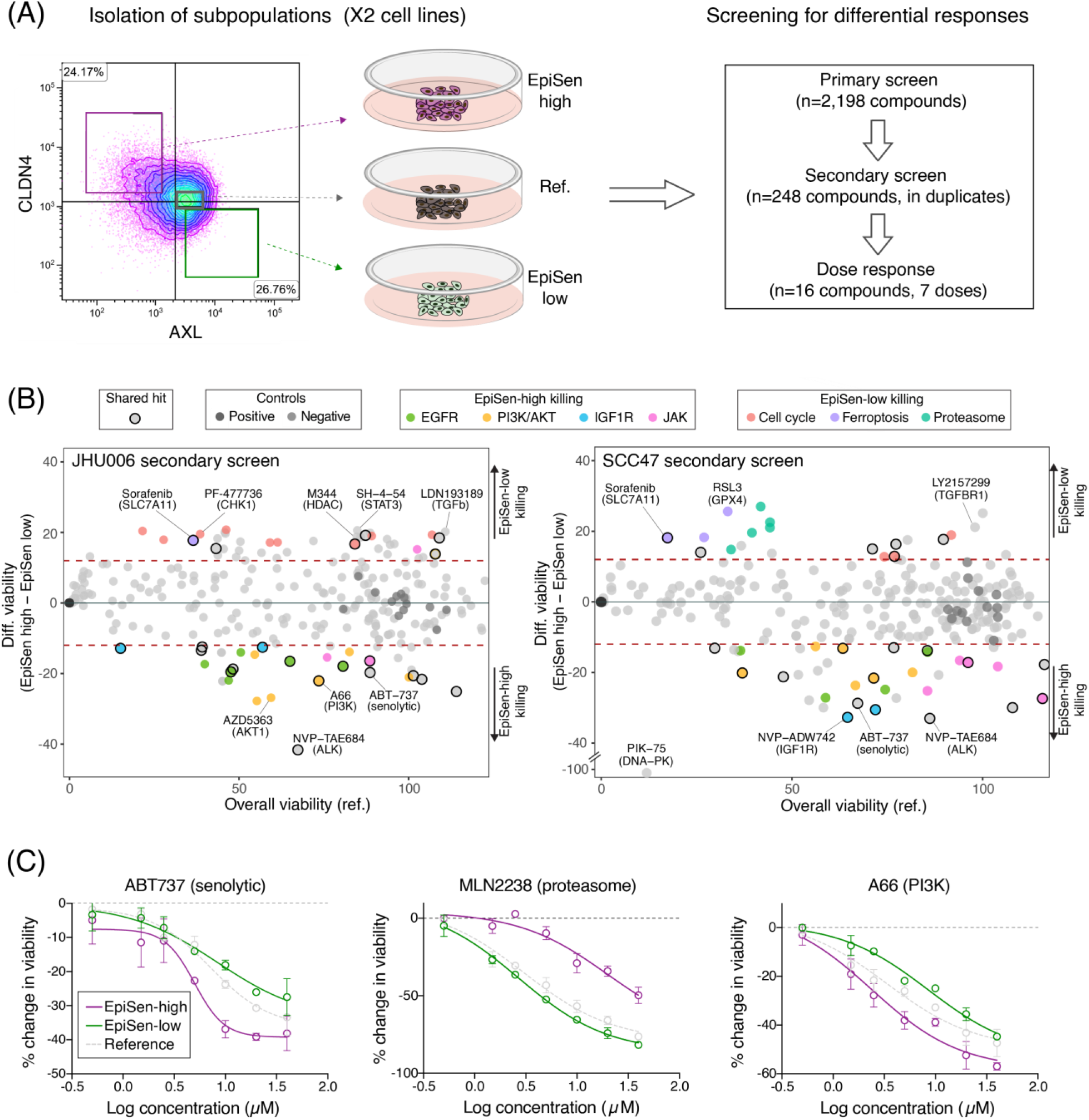
Co-existing cellular states differ in drug sensitivity. **(A)** Experimental scheme for drug screening: three subpopulation were isolated by FACS and subjected to primary screen, secondary screen and dose response analysis of selected hits. **(B)** Viability of the reference population (X-axis) and differential viability of the EpiSen-high vs. EpiSen-low populations (Y-axis) upon treatment with 248 compounds tested in the secondary screen, in JHU006 (left) and SCC47 (right). Dotted lines represent thresholds for differential sensitivity. Selected hits and controls are colored as specified in the top legends. **(C)** Dose response curves of three selected compounds in SCC47, presented by the change in viability relative to vehicle controls. Error bars represent standard deviation.

We initially screened 2,198 bioactive compounds using a CTG-based viability assay (**fig. S8A-C).** We identified 200 compounds (9%) as potential “hits”, defined by differential killing of EpiSen-high and EpiSen-low subpopulations in at least one cell line. There was a significant overlap among compounds that preferentially killed EpiSen-high cells in the two cell lines (*P*=0.006, hypergeometric test). Next, these putative hits and an additional group of compounds that killed both populations in the primary screen (*n*=248) were selected for a secondary screen performed in duplicate in each cell line (Fig. 6B).

The secondary screen identified 113 compounds with differential killing of the subpopulations in at least one cell line. Of the hits preferentially killing the EpiSen-high cells, 15 were shared among both cell lines, representing 41% and 45% of all the corresponding hits in JHU006 and SCC47, respectively. This fraction of consistent hits further increases to 71.4% (for both cell lines) when considering the targets of compounds rather than the exact compounds, highlighting consistent vulnerabilities of EpiSen-high cells. Finally, fourteen compounds with differential sensitivities, including five that were shared between cell lines and nine that were specific to one cell line, were analyzed by a full dose response (Fig. 6C, **fig. S8D**). All five of the shared compounds, and five of the nine cell line-specific compounds (56%), displayed significant differential sensitivity as in the secondary screen (*P<*0.05, paired t-test), supporting the consistency between cell lines as a measure of robustness.

As expected, EpiSen-high cells were more sensitive to the senolytic compound ABT-737 (Yosef et al., 2016). Additional EpiSen-high sensitivities included multiple inhibitors of EGFR, AKT, PI3K, DNA-PK, IGF1R, and JAK (Fig. 6B). Several of these targets (DNA-PK, IGF1R and AKT) converge on repair of double-strand breaks as part of the DNA repair machinery (Bozulic et al., 2008; Wong et al., 2009). Together with the observation that hydrogen peroxide induces the expression of EpiSen genes (Fig. 5E), these results reinforce the role of DNA damage as a potential inducer of this RHP. The PI3K/AKT axis is hyper-activated in HNSCC, and resistance to PI3K inhibition in HNSCC is AXL-dependent (Elkabets et al., 2015). Accordingly, EpiSen-high cells (which are defined by low AXL expression) were more sensitive to inhibitors of PI3K and AKT, as well as those of EGFR and IGF1R that signal via the PI3K/AKT axis.

EpiSen-low cells were more sensitive to inhibitors of cell cycle regulators (CDKs, CHK1 and topoisomerase), consistent with their increased proliferation. EpiSen-low sensitivities also included multiple inhibitors of the proteasome and compounds that induce cell death by sensitizing cells to ferroptosis (the GPX4 inhibitor RSL3, Erastin, and the SLC7A11 inhibitor Sorafenib) (Fig. 6B). Recent work demonstrated that mesenchymal cells are particularly sensitive to ferroptosis-inducing compounds (Hangauer et al., 2017; Viswanathan et al., 2017). Thus, some of these vulnerabilities may reflect an increased EMT of EpiSen-low cells due to the inverse correlation between EpiSen and EMT-II. Taken together, these results suggest that EpiSen-high and EpiSen-low cells are associated with differential vulnerabilities that may be consistent across distinct cellular contexts.

## Discussion

Despite widespread appreciation of the importance of tumor cell heterogeneity and emerging single cell profiling methods, deciphering the function and mechanisms underlying the observed heterogeneity requires follow up experiments that are not feasible with clinical samples. Identifying model systems that recapitulate cellular diversity within tumors (or aspects thereof) is a critical step towards this goal. One approach is to continue developing more realistic models of tumors, such as humanized mouse models and three-dimensional organoid cultures, which benefit from complex tumor-like microenvironment. In parallel, it is important to more rationally utilize existing *in vitro* cultures, where hundreds of models can be screened in parallel, experiments are considerably easier, and there is a large body of legacy research. The relevance of such an approach depends on whether or not it is possible to recapitulate important aspects of intra-tumoral heterogeneity in such simple model systems. While cell lines are often criticized and considered inadequate, we do not really know the extent of diversity of cellular states within them, and whether it relates to the diversity observed within tumors. Recapitulation of *in vivo* diversity in cell lines also highlights cases where plasticity is at least partially intrinsic to the cancer cells, and thereby observed in the absence of a native microenvironment.

Our multiplexing strategy allowed us to effectively profile ~200 cancer cell lines by scRNA-seq. While individual cell lines may be influenced by other cell lines in their pool, previous work demonstrated that cell lines in these pools retain their differential drug sensitivities (Yu et al., 2016). Moreover, we focused exclusively on the variability between cells from the *same* cell line (rather than on the global expression pattern of each cell line), which is less sensitive to pooling and batch effects (**fig. S3A**) and is indeed consistent across different pools (**fig. S3B-C**) as well as between our scRNA-seq profiles and our follow up experiments with two HNSCC cell lines.

The main component of expression heterogeneity in cancer cell lines was continuous, with only a few cases of discrete subpopulations within the same cell line. Such gradual patterns contrast with the discrete nature of genetic subclones, suggesting at least a partial decoupling between genetic heterogeneity and the nongenetic heterogeneity that might underlie continuous patterns. Consistent with this hypothesis, we found only limited association between genetic subclones (by inferred CNAs) and expression heterogeneity, particularly when considering continuous patterns. Moreover, follow up experiments directly demonstrated the dynamic plasticity of the EpiSen program. Thus, cancer cells may harbor variability through two largely distinct processes of genetic and nongenetic mechanisms, both of which may contribute to drug resistance and tumor progression, with limited interaction between them. We further speculate that by focusing on recurrent patterns of heterogeneity our analysis highlights nongenetic plasticity, as this form of variability tends to be shared across cell lines, while genetic forms of variability may be more unique to each cell line.

Nine of the 12 recurrent programs we identified (including cell cycle RHPs) were also observed in tumors *in vivo*, indicating that they are partially retained even in the absence of an *in vivo* microenvironment, and may primarily reflect cell-intrinsic plasticity. As each RHP was observed in a subset of cell lines, we leveraged the extensive cell line annotations to identify associations between RHPs and genetic backgrounds (p53, MRE11A and NOTCH4), lineages (melanoma and HNSCC), and drug sensitivities (*e.g.*, Nutlin-3a). Moreover, we further prioritized particular cell lines that most reliably mirror ITH patterns for follow up experiments. This is an important deviation from the traditional focus on historical cell lines that are easy to grow, and provides a resource for custom prioritization of cell lines based on their heterogeneity.

Careful examination of the programs *in vivo* and *in vitro* highlights the divergence from their developmental “normal” counterparts. Both EMT and senescence are associated with precise phenotypes and well-defined regulators during development and wound healing, yet in the context of tumors and cancer cell lines, we observe only partial phenotypes and limited dependence on these regulators. The EMT-like profiles we observe include many EMT-related genes and are associated with increased migration, but do not involve EMT hallmarks such as the loss of epithelial markers, a drastic change in morphology, and expression of most EMT transcription factors. Similarly, EpiSen-high cells resemble the senescence response of keratinocytes and lung bronchial cells, are associated with reduced proliferation, and possess markers of SASP, yet they retain some proliferative capacity, harbor p53 mutations and do not express high levels of p16 and p21. This is consistent with studies showing evidence for incomplete and reversible senescence programs in cancer. Low levels of p16 at induction of senescence confers cell cycle re-entry upon p53 inactivation or RAS expression (Beausejour et al., 2003), a program coined “light senescence” (Lee and Schmitt, 2019) and likewise, loss of Rb in senescent cells leads to renewed proliferation (Sage et al., 2003). We hypothesize that cancer cells often activate partial or distorted programs, possibly not through the canonical developmental mechanisms, and in a context-dependent manner. This could contribute to the difficulties in resolving long-standing debates in the cancer field about the role of EMT and senescence, which are often evaluated through the activity of developmental regulators and markers that may fail to detect certain partial programs. Comprehensive single cell profiling helps detect such partial programs that vary in their magnitude across the cells.

A partial and reversible epithelial senescence (EpiSen) state in tumors could have several implications. First, EpiSen may continue to restrict the growth rate of cancer cells even in established tumors and cell lines. Second, EpiSen cells may be particularly resistant to certain treatments (Fig. 6), such as chemotherapies that target proliferating cells, and thereby may facilitate tumor recurrence. Third, EpiSen could function as a DNA repair program, allowing damaged cells to evade apoptosis and/or ferroptosis. This is consistent with the induction of EpiSen by hydrogen peroxide, the sensitivity of EpiSen-high cells to inhibitors of DNA repair, and the functions of core EpiSen genes in oxidative stress, including the oxidant scavengers S100A8/9, the H202 transporter AQP3, and LCN2, a secreted factor that increases ROS levels (Kagoya et al., 2014). Fourth, EpiSen could remodel the tumor microenvironment, influencing cancer, stromal and immune cells through its abundance of secreted factors, including S100A8/A9, SAA1/2, SLPI, CXCL1, and LCN2. While SASP is best characterized in fibroblasts, here we describe a distinct “EpiSASP” whose function will be investigated by follow up studies. Lastly, EpiSen cells re-entering the cell cycle may confer unique properties such as increased tumor initiation capacity (Milanovic et al., 2018).

ITH has long been recognized as a potential cause of therapeutic failure. Our analysis supports this notion by demonstrating that subpopulations of cells from the same cell line are associated with distinct drug sensitivities. We ensured the robustness of these results by three approaches: (1) Comparing pairs of subpopulations from the same cell line, to control for cell line-specific drug sensitivities; (2) Focusing on effects that were consistent across a HPV+ and a HPV-HNSCC cell lines that differ in many regards; and (3) Validating selected hits with a secondary screen as well as a full dose-response analysis. The differential responses we identified included EGFR inhibitors, which are routinely used in the treatment of HNSCC patients, underscoring the potential clinical relevance of our observations. Our results suggest that EGFR inhibitors (as well as other inhibitors) preferentially eliminate subsets of EpiSen-high cells, providing a rationale for their combination with chemotherapies that target the more proliferative EpiSen-low subpopulations.

In conclusion, we described here the landscape of diversity across ~200 cell lines of various cancer types, generating a dataset that would be widely useful for the cancer research community. In analyzing this extensive dataset, we found multiple recurrent programs of heterogeneity that recapitulate ITH, and are associated with continuous nongenetic plasticity. Follow up analysis of one such program demonstrated its dynamics, regulation and vulnerabilities. Further studies of tumors and of the model systems prioritized through this data will provide a better understanding of ITH, which is currently a main barrier for successful cancer therapies.

## Methods

### Cell line pools

We obtained eight previously generated pools of cell lines (Yu et al., 2016), each containing 24-27 cell lines from diverse cancer types. Cell lines were combined to pools based on growth rates (doubling time), in order to ensure comparable representation over the short-term culturing. Each pool was thawed and cultured in RPMI medium supplemented with 10% fetal bovine serum for 3 days before scRNA-seq. A ninth custom pool was generated by freshly pooling eight additional head and neck cell lines immediately prior to scRNA-seq (JHU006, JHU011, JHU029, SCC9, SCC90, SCC25, UM-SCC47, 93VU-147T).

### Individual cell line cultures

Human HNSCC cell lines (laryngeal: JHU006, JHU011, JHU029; oropharyngeal: UM-SCC47, SCC9, SCC90, SCC25, 93-VU-147T) were provided by Dr. James Rocco after confirmation by short tandem repeat analysis. The laryngeal cell lines were grown in RPMI 1640 media (Biological Industries, Kibbutz Beit HaEmek). The oropharyngeal cell lines were grown in a 3:1 mixture of Ham’s F12:DMEM (Biological Industries, Kibbutz Beit HaEmek). All growth media for HNSCC cell lines was supplemented with 10% fetal bovine serum (Biological Industries, Kibbutz Beit HaEmek), 1x penicillin-streptomycin and 1x L-glutamine (Biological Industries, Kibbutz Beit HaEmek).

Human primary bronchial epithelial cells (PCS-300-010) were acquired from ATCC and grown in Airway Epithelial Cell Basal Medium (ATCC PCS-300-030) supplemented with the bronchial epithelial cell growth kit (ATCC PCS-300-040) consisting of HSA 500 µg/mL, linoleic acid 0.6 µM, lecithin 0.6 µg/mL, glutamine 6 mM Extract P 0.4%, Epinephrine 1.0 µM, Transferrin 5 µg/ml, T3 10 nM, Hydrocortisone 0.1 µg/ml, EGF 5 ng/mL, and Insulin 5 µg/mL. All cell lines screened negative for mycoplasma by the EZ-PCR mycoplasma detection kit (Biological Industries, Kibbutz Beit HaEmek).

### Droplet-based scRNA-seq

ScRNA-seq libraries were generated using the 10X Genomics Chromium Single Cell 3’ Kit v2 and the 10x Chromium Controller (10x Genomics) according to the 10X Single Cell 3’ v2 protocol. Briefly, for each pool, we generated a single cell suspension in 0.04% PBS-BSA (≥95% viability) and loaded approximately 10,500 single cells to the Chromium Controller with a targeted recovery of 6,000 cells. Single cells, reagents and single gel beads containing barcoded oligonucleotides were encapsulated into nanoliter-sized droplets and subjected to reverse transcription. Droplets were broken and the barcoded cDNAs were purified with DynaBeads and amplified by 12 cycles of PCR (98°C for 45 s; [98°C for 20 s, 67°C for 30 s, 72°C for 1 min] x 12; 72°C for 1 min). The amplified cDNA was fragmented, end-repaired, ligated with index adaptors, and size-selected with clean-ups between each step using the SPRIselect Reagent Kit (Beckman Coulter). Quality control of the resulting barcoded libraries was performed with the Agilent TapeStation and by PCR with primers specific to the P5 and P7 sequence (NEBNext Library Quant Kit for Illumina, New England Biolabs).

### Bulk RNA-seq by SMART-Seq2

The SMART-Seq2 protocol (Picelli et al. 2014) was adapted with several modifications, as described below, to generate libraries for bulk RNA-Seq of individual cell lines included in the HNSCC custom pool (see ‘Cell line assignment’) and for profiling of bronchial primary cells following senescence induction with etoposide (**fig. S4E**) (Rauner et al., 2018). Between 100-200 cells were resuspended in lysis buffer containing Triton 0.2% and RNAase inhibitor, or pelleted, washed with PBS and resuspended in the lysis buffer. For library generation, the plate was incubated at 72°C for 3 min and reverse transcription was performed as described in the SMART-Seq2 protocol (Picelli et al., 2014) followed by cDNA pre-amplification for 17 cycles. Following 1X Agencourt Ampure XP beads cleanup (Beckman Coulter), 200 pg of amplified DNA underwent tagmentation and final amplification adding unique Illumina barcodes for 12 cycles (Nextera XT Library Prep kit, Illumina). Libraries were quantified by Qubit and TapeStation (Agilent) and pooled prior to sequencing. Sequencing was performed using the Illumina Nextseq 75 cycle high output kit (single read, dual index).

### Bulk RNA-seq by MARS-Seq

A bulk adaptation of the MARS-Seq protocol (Keren-Shaul et al., 2019) was used to generate RNA-seq libraries for validation of isolation of selected subpopulations by FACS (Fig. 5B) and for expression profiling of HNSCC cell lines following perturbations (Fig. 5E, **fig. S7D**). RNA was isolated from sorted and unsorted cells using the Quick RNA Microprep kit (Zymo Research). Briefly, 50 ng of input RNA from each sample was barcoded during reverse transcription and pooled. Following Agencourt Ampure XP bead cleanup (Beckman Coulter), the pooled samples underwent second strand synthesis and were linearly amplified by T7 in vitro transcription. The resulting RNA was fragmented and converted into a sequencing-ready library by tagging the samples with Illumina sequences during ligation, RT, and PCR. Libraries were quantified by Qubit and TapeStation as well as by qPCR for GAPDH, as previously described (Keren-Shaul et al., 2019). Sequencing was done with the Illumina Nextseq 75 cycle high output kit (paired-end sequencing).

### Sequencing

Final 3’ scRNA-Seq libraries were diluted to 4 nM, denatured, and further diluted to a final concentration of 2.8 pM for sequencing with the following sequencing parameters: Read 1: 26 cycles, i7 index: 8 cycles, i5 index: 0 cycles, Read 2: 58 cycles. Pooled bulk MARS-Seq libraries were diluted to 4 nM, denatured, further diluted to 2 pM and sequenced with the following parameters: Read 1: 75 bp, Read 2: 15 bp, no indices. Bulk Smart-Seq2 libraries were diluted to 1.6 pm and sequenced with parameters: Read 1: 76bp, i7: 8 cycles, i5:8 cycles. All sequencing was performed with the NextSeq500 (Illumina) using the NextSeq 75bp High Output Kit (Illumina), either at the Broad Institute or at the Weizmann Institute Life Science Core Facility.

### Flow cytometry and sorting of cell lines

Sorting of JHU006 and UM-SCC47 cells was performed on a BD FACS Melody using the following antibodies: anti-human AXL-PECy7 (eBioscience) at 1:300, anti-human Claudin-4-APC (Miltenyi) at 1:200, and anti-human ITGA6/CD49f-APC eBioscience) at 1:200. Gating of positive and negative cells was defined by the unstained control.

For EpiSen program dynamics experiments, 200,000 cells of each subpopulation (EpiSen-high: AXL^−^CLDN4^+^; EpiSen-low: AXL^+^CLDN4^−^; control sort: all single cells) were sorted and reanalyzed by FACS immediately post-sorting and at days 7 and 14. Final analysis was performed using Kaluza Analysis Software v2.1 (Beckman Coulter). The experiment was performed three times independently. Successful isolation of the EpiSen subpopulation by AXL and Claudin-4 was validated by bulk RNA-Seq of sorted cells as described in the ‘Bulk RNA-Seq by MARS-Seq’ section above. Similarly, the EMT-II subpopulation was isolated by AXL^+^ITGA6^+^ (EMT-II-high cells) and AXL^−^ITGA6^−^ (EMT-II low cells).

### Cell cycle analysis by propidium iodide (PI) DNA staining

The EpiSen-high (AXL^−^CLDN4^+^) and EpiSen-low (AXL^+^CLDN4^−^) subpopulations were isolated from the JHU006 and UM-SCC47 cell lines as described above. Cells were sorted into ice cold PBS and fixed by adding the cell suspension dropwise to 70% ethanol while vortexing. Fixed cells were stored at 4°C. Following 2x PBS washes, cells were resuspended and incubated in PI/Triton-X-100 staining solution consisting of 0.1% Triton-X-100 (Sigma), 0.2mg/ml DNAse-free RNAse A (Sigma), and 0.04 mg/ml of 500ug/ml PI (Sigma) in PBS at 20°C for 30 min. Singlets were gated by FSC-W vs. FSC-A (PI) and cell cycle analysis was performed based on the PI signal histogram. The experiment was performed three times independently.

### Cytokine treatments and perturbations of HNSCC cell lines

JHU006 and SCC47 cells were seeded at 50,000 cells/well in 24 well plates in their standard media and treated in duplicate with drug/cytokine or vehicle (0.1% DMSO with 1ug/mL BSA) 24 hours after seeding. Cells were harvested 24 hours after treatment and RNA was isolated by the Quick RNA Microprep kit (Zymo Research) for bulk expression profiling using the MARS-Seq protocol (above). Treatments included 10 μM all-trans-retinoic acid (ATRA) (Sigma), 25 ng/ml IFN-γ (Peprotech), 25 ng/ml TNF-α (Peprotech), 25 ng/ml PDGF-BB (Miltenyi), 10 nM etoposide (Sigma), 10ng/ml TGF-β1 (Peprotech), 10ng/ml TGF-β3 (Peprotech), 10 μM cisplatin (Sigma), 200 μM hydrogen peroxide (Sigma), 25 ng/ml IL-8/CXCL8 (Peprotech), 25 ng/ml GAS6 (Sino Biological), 50 ng/ml S100A8/A9 (Sino Biological), and 500 μM desferoxamine (DFO) (Sigma).

### Senescence induction by etoposide and SA-β-gal staining

Primary bronchial cells were seeded at 50,000 cells/well in 24 well plates and treated with 5-7.5 μM etoposide (Sigma) 24 hours later to induce senescence. After 48 hours, media was replaced and on day 9 etoposide-treated cells and untreated controls were fixed with 0.5% glutaraldehyde solution in PBS pH 7.4 and incubated with X-gal staining solution (0.2M K_3_Fe(CN)_6_, 0.2M K_4_Fe(CN)_6_ 3H_2_O, and 40X X-Gal stock diluted in PBS/MgCl_2_) for 6 hours protected from light. X-Gal stock consists of 40 mg/ml X Gal (Roche #745740) in N,N-dimethylformamide (Sigma D-4254). Following PBS washes, stained cells were covered with 80% glycerol prior to imaging.

### Gap closure migration assay

Following sorting by AXL and ITGA6 to isolate the EMT-II-high and EMT-II-low subpopulations as described above, 75,000 cells of each population (EMT-II-high cells, EMT-II-low cells, unsorted cells, and unsorted cells treated with 10 ng/µl TGF-β3 (Peprotech)) were resuspended in 70 µL RPMI and loaded per well containing a wound healing assay culture insert (Ibidi). After 24 hour incubation, the inserts were removed to generate the gap and cells were imaged at 0, 6, 12, 24, and 48 hours. The experiment was performed independently three times.

### Drug screening - viability assay

The Selleck Bioactive Compound Library (Selleck Chemicals) as well as DMSO-only controls and straurosporine positive (killing) controls were dispensed into 384-well plates with an Echo 550 liquid handler (Labcyte). Drug concentration was 10 µM for the primary screen and 1 µM or 10 µM for the secondary screen (performed in duplicate) depending on hit category. Purity of compounds selected for follow up by dose response was confirmed by LC/MS (data not shown). For the dose response curves, a seven-point, two-fold dilution series with an upper limit of 40 µM was tested in duplicate. EpiSen-high (AXL^−^CLDN4^+^), EpiSen-low (AXL^+^CLDN4^−^), and a third neutral reference population were sorted from JHU006 and SCC47 cell lines, as described above, and sorted subpopulations were seeded into the compound-treated plates in their standard media at a concentration of 15,000 cells/ml (750 cells/well) with a Combi Multi-drop (ThermoFisher). Plates were incubated at 37°C for 48 hours following sorting and compound treatment and cell viability was determined based on luminescence following addition of CellTiter-Glo (Promega) according to the manufacturer’s instructions. Luminescence was measured on a BMG Pherastar plate reader. Data was normalized in Genedata Screener where DMSO (vehicle) is defined as neutral control (i.e. 100% viability) and samples without cells are inhibitor control (i.e. 0% viability). Compound-centric data was visualized in CDD Vault from Collaborative Drug Discovery (Burligame, CA).

### Processing of scRNA-seq data

Cell barcode filtering, alignment of reads and UMI counting were performed using CellRanger 3.0.1 (10x Genomics). Expression levels were quantified as E_i,j_ = log_2_(1 + CPM_i,j_/10), where CPM_i,j_ refers to 10^6^*UMI_i,j_/sum[UMI_1..n,j_], for gene *i* in sample *j*, with *n* being the total number of analyzed genes. The average number of UMIs detected per cell was less than 100,000, thus CPM values were divided by 10, to avoid inflating the differences between detected (E_i,j_ > 0) and non-detected (E_i,j_ = 0) genes, as previously described (Tirosh et al., 2016b). For each cell, we quantified the number of detected genes as a proxy for sample quality. We conservatively retained cells with a number of detected genes ranging from 2,000 to 9,000. When analyzing cell lines individually, we only considered genes expressed at high or intermediate levels (E_i,j_ > 3.5) in at least 2% of cells, yielding an average of 6,758 genes analyzed per cell line. Values were then centered per cell line to define relative expression values, by subtracting the average expression of each gene *i* across all k cells: Er_i,j_=*E*_*i,j*_-*average*[*E*_*i*,1...*k*_]. When analyzing cell lines collectively, we selected the top 7,000 expressed genes across all cell lines, resulting in a minimum average expression of 12 CPM. Values were centered by subtracting the average expression across all 53,513 cells analyzed: *ER*_*i,j*_ = *E*_*i,j*_-*average*[*E*_*i*,1…53,513_].

### Processing of bulk RNA-seq data

Reads were aligned to the GHCh38/hg38 human genome using Bowtie and expression values were quantified using RSEM. Data are presented as E_i,j_ = log_2_[(TPM_i,j_) + 1], where TPM_i,j_ refers to transcript-per-million for gene *i* in sample *j*, as calculated by RSEM (Li and Dewey, 2011).

### Cell line assignment

We used both expression-based and SNP-based methods to assign cells to cell lines. In each of these methods, we compared the single cells to external bulk data of the corresponding cell lines and then either assigned the cells to the most similar cell line or excluded them as potential doublets or low-quality cells. Only cells whose assignments were also consistent between both methods were retained for further analysis. Bulk RNA-seq data was obtained from the DepMap portal (https://depmap.org/; 18q3 data release) (Ghandi et al., 2019) for cell lines in the eight CCLE-based pools and was generated as described above for the eight HNSCC cell lines in the custom pool.

For SNP based classification, for each cell we determined the cell line from the pool whose SNP profile (based on bulk RNA-seq data) best matched the observed reference and alternate allele reads across a panel of SNP sites, similar to the Demuxlet method (Kang et al., 2018). Specifically, we used a logistic regression model for each cell, where the probability of a read at SNP site *i* being from the alternate allele is modeled as *P*_*i*_ = *σ*(*β*_*0*_+*β*_*j*\*_*X*_*i,j*_), where *σ* is the logistic function, *X*_*i,j*_ is the allelic fraction of cell line *j* at site *i* (estimated from bulk RNA-seq data), and *β*_*j*_ care parameters estimated for each single cell and reference cell line by maximizing the data likelihood under a binomial model. Models were fit using the R package glmnet (Friedman et al., 2010), and the cell line whose SNP profile produced the highest likelihood under this model was selected. Goodness-of-fit was quantified by the model deviance relative to the null-model deviance. We used a reference panel of 100,000 SNPs that were frequently detected across a panel of 200 cancer cell lines (based on bulk RNA-seq data), and that were detected in the scRNA-seq data from the same cell lines. For expression-based classification, we used a broadly similar approach. First, we subsetted the gene expression data to genes that were expressed in at least half of all cells or had a maximal expression (measured by the 98^th^ percentile of that gene’s expression across all cells) greater than 3 log_2_(CPM). Next, we used the Rtsne R package to estimate, for each pool, 3 t-SNE embedding dimensions for each cell, computed based on 50 principle components, using a perplexity parameter of 30, and a theta value of 0.2 (using the first three principle components for initialization). We applied a local ‘smoothing’ to the normalized and centered single-cell expression profiles (*ER*), using a Gaussian kernel applied to the cell-cell distances in the t-SNE embedding space. The Gaussian kernel bandwidth was set using the method ‘sigest’ from the R package kernlab. Finally, for each cell, we identified the reference cell line from the pool with the most similar bulk RNA-seq gene expression profile (*ER*, and Pearson correlation similarity).

Detection of putative ‘doublets’, where two or more cells are labeled with the same barcode during droplet-based library preparation, was performed based on the SNP data, using the same generalized linear modeling approach to identify a mixture of two reference cell lines whose combined SNP profiles best explained the SNP data from a given cell. To efficiently estimate the best-fitting reference cell line pair we used a Lasso-regularized generalized linear model. After determining the best-fitting ‘singlet’ and ‘doublet’ models for each putative cell, we then determined whether each putative cell was a singlet, doublet, or a ‘low-quality’ cell based on several statistics. To identify low quality cells we took the maximum of the deviance explained by the singlet model and the deviance explained by the doublet model. We observed that the maximum deviances formed a bimodal distribution. We thus used the local minimum between the two distributions as a threshold and classified all cells with a maximum deviance below this threshold as ‘low quality’. To separate putative doublets from singlets, we then fit a two-component Gaussian mixture model using three variables: (1) the amount of deviance explained by the singlet model; (2) the (log-transformed) deviance-improvement of the doublet model over the singlet model; and (3) the fraction of genes detected in that cell. Cells with a probability greater than 0.75 of belonging to the cluster with higher average doublet improvement were classified as doublets.

### Systematic characterization of transcriptional heterogeneity

For each of the 198 cell lines passing QC, we applied two distinct approaches to identify discrete and continuous patterns of expression heterogeneity. First, to identify discrete (highly distinct) subpopulations within cell lines, we used nonlinear dimensionality reduction (t-SNE) followed by density-based clustering (DBSCAN), which assumes that clusters are contiguous regions with high density of cells. T-SNE was applied to each cell line individually using relative expression values (*Er*) and perplexity of 30. To identify dense regions, DBSCAN classifies each point according to a minimum points (minPts) threshold, defined as the minimum number of neighbors within a user-defined radius (eps) around core points. To optimize this parameter selection, we tested the ability of DBSCAN to correctly distinguish cells from two distinct cell lines. We combined cells from two different cell lines and tested the classification accuracy of DBSCAN using different eps (0.6 – 3) and minPts thresholds (5 and 10). DBSCAN classification was evaluated using Fisher’s exact test and considered correct if p < 0.001. We repeated this procedure 1,000 times and in each iteration we randomly selected the cell lines, the total number of cells (56-1,990) and the proportion of cells selected from each cell line (2-98% of total). The parameter combination yielding the highest rate of correct classification (eps = 1.8, minPts = 5) was used in further analyses. We also applied DBSCAN with additional, less stringent eps values (1.2 and 1.5) to show the robustness of the results. To define gene signatures that characterize the discrete subpopulations identified, we compared gene expression of cells in a given cluster to all other cells within the same cell line using a t-test. Genes with fold-change>=2 and *P*<0.001 were selected and the top 50 (by fold change) were defined as the gene signature. Clusters containing above 90% of the cells of a given cell line were excluded from this analysis.

Second, we analyzed each cell line separately using NMF to identify both discrete and continuous programs of expression heterogeneity. NMF was applied to the relative expression values (*Er*), by transforming all negative values to zero, as previously described (Puram et al., 2017). We performed NMF with the number of factors *k* ranging from 6-9, and initially defined expression programs as the top 50 genes (by NMF score) for each *k*. For each cell line, we sought robust expression programs by selecting those with an overlap of at least 70% (35 out of 50 genes) with a program obtained using a different value of *k*. To avoid redundancies, from each set of overlapping programs from one cell line, we only kept one program, selected based on having the highest overlap with a NMF program identified in another cell line. To determine which programs reflected discrete patterns of expression variability, for each cell line we compared the classification of cells by DBSCAN and the assignment of cells to the highest-expressed NMF program using Fisher’s exact test (p < 0.001 were considered significant). The association between programs and technical artifacts was inspected by calculating, for each cell line, the Pearson correlation coefficient between the number of genes detected in a cell (complexity) and the respective NMF scores. This approach identified a cluster of NMF programs (based on 50 minus the number of overlapping genes across expression programs) that share negative correlations with complexity. This cluster appeared to reflect technical artifacts (based also on manual inspection and inclusion of many mitochondrial genes and pseudogenes) and was excluded from further analysis.

In order to identify recurrently heterogenous expression programs (RHPs) across cell lines, we compared expression programs derived from DBSCAN and NMF, separately, by hierarchical clustering, using 50 minus the number of overlapping genes as a distance metric. Given the high number of NMF programs, clustering was restricted to programs with at least a minimum overlap of 20% (10 out of 50 genes) with a program observed in another cell line. Twelve clusters were defined by manual inspection of the hierarchical clustering results. For each cluster of program, a RHP was then defined as all genes included in at least 25% of the constituent programs. We then used the hypergeometric test to assess the enrichment of RHP signatures with H and C5:BP gene-sets from MSigDB (Liberzon, 2014), and p < 0.001 was considered significant.

### Pool effect analysis

To evaluate the impact of the pooling procedure on the cell lines’ expression we used three different approaches. First, we determined the proportion of the similarity dependent on the pool of origin in either the mean expression profiles of cell lines or in expression heterogeneity patterns (by NMF). To this end, we calculated pairwise correlations between mean expression profiles of the cells in each cell line, and used one-way ANOVA to compute the proportion of the total variance (η^2^) explained by whether or not the cell lines were in the same CCLE pool. This same procedure was applied to evaluate the pairwise correlations between NMF programs across gene scores. Second, we inspected the distribution of each RHP across all CCLE pools, to evaluate whether there were pool-specific biases. Third, we applied NMF (with 6 factors) to scRNA-seq data of the HNSCC cell line SCC25, which was profiled in two different conditions: as part of the CCLE pool ID #19 and as part of the custom HNSCC pool. We compared programs by hierarchical clustering using one minus Pearson correlation coefficient across NMF gene scores as a distance metric.

### Defining program scores in each cell

Program scores were calculated for each cell individually in order to evaluate the degree to which they express a given RHP. Cells with higher complexity (defined as larger number of genes detected) would be expected to have higher cell scores for any gene-set. To account for this effect, for each gene-set analyzed, we created a control gene-set to be used in the calculation of a normalization factor, as previously described (Tirosh et al., 2016b). Control gene-sets are selected in a way that ensures a similar distribution of expression levels to that of the input gene-set. First, all genes analyzed are ordered by mean expression across all cell lines and partitioned into 75 bins. Next, for each gene in the given gene-set, we randomly select 100 genes from the same expression bin. Finally, given an input set of genes (*G*_*j*_), we defined a score, *SC*_*j*_(*i*), for each cell *i*, as the mean relative expression of the genes in *G*_*j*_. We then calculate a similar cell score for the respective control gene-set and subtract it from the initial cell scores: 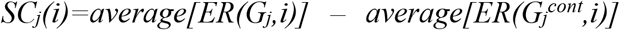.

### *Comparison of* in vitro *and* in vivo *programs*

In vivo *programs of expression heterogeneity were previously defined (Puram et al., 2017; Tirosh et al., 2016a) or generated using published scRNA-seq data (Chung et al., 2017; Lambrechts et al., 2018) with the NMF-based strategy described above applied to the malignant cells in each dataset separately. For each* in vivo *program we calculated the maximum similarity (Jaccard index) and maximum single cell score (*SC*) correlation with* in vitro *programs. Correlations were calculated using the cell line harboring the respective* in vitro *program. Next, we identified the overall maximum similarity and maximum correlation obtained by permuting each* in vitro *program 100 times. To generate permuted gene-sets we first ordered all genes by mean expression across all cell lines and partitioned them into 75 bins. Each gene in a given gene-set was then replaced by a randomly select gene from the same expression bin.* In vivo *programs presenting both maximum similarity and maximum correlation above the permutation threshold were selected. Among selected programs we focused on the top 10 with highest correlation, and manually identified 5 main groups: melanoma MITF, melanoma AXL, HNSCC epithelial differentiation, HNSCC pEMT and melanoma/HNSCC stress. We defined which* in vitro *RHP most closely recapitulate the 5 selected* in vivo *programs based on the average similarity and correlation of the constituent programs*.

### Computational cell cycle analysis

Scoring cells for the G1/S and G2/M RHPs reveals a circle-like structure, reflecting different phases of the cell cycle (**fig. S3C**). This pattern recurs across cell lines, and was also previously described for different human cancers and mouse hematopoietic stem cells (Kowalczyk et al., 2015; Tirosh et al., 2016a; Tirosh et al., 2016b). Since these patterns are continuous, borders between cell cycle phases were defined conservatively by manual inspection into four patterns (**fig. S3C**): non-cycling (SC_G1/S_ < −0.75 and SC_G2/M_ < −0.5), G1 (SC_G1/S_ > −0.5 and SC_G2/M_ < 0), S (SC_G1/S_ > 0.25 and SC_G2/M_ > 0), and G2/M cycling (SC_G1/S_ < 0.25 and SC_G2/M_ > 0.5).

### Defining program variability in each cell line

To evaluate the degree of heterogeneity of RHPs in each cell line, we examined the variability of cell scores. First, given a program *j* and cell line *i*, we defined program variability, *PV*_*j*_(*i*), by first ranking cells according to the program score (*SC*_*j*_) and comparing the average signal between the top and bottom 10% cells: *PV(j)=average[SC*_*j*_(*top10%*)] − *average[SC*_*j*_(*bottom10%*)]. Next, to control for the potential association between mean and variability of program scores, we applied a local polynomial regression with a smoothing span of 0.8 to infer the relationship between program variability and mean in each cell line, and used the residuals of the model as the corrected program variability score: *PVc(j) = PV(j) - RG(mean(SCj))*, where RG represents the local regression model for PV based on the average program scores, SC.

### Association between program variability and mutations or drug responses

Mutation calls (coding region, germline filtered) were downloaded from the CCLE portal (https://portals.broadinstitute.org/ccle), and drug response data (CTRP v2, area under the curve, AUC) were downloaded from the CTD^2^ portal (https://ocg.cancer.gov/programs/ctd2/data-portal) (Basu et al., 2013). We restricted the analysis to non-silent mutations and compounds tested in at least 160 out of the 198 cell lines analyzed. We compared program variability scores of mutated and wild-type cell lines using two-sided t-test. The association between drug sensitivity (1-AUC) and program variability scores was assessed using multiple linear regression, with cancer type as a covariate: *Yd* ∼ *cancer type* + *PV*_*j*_, for drug *d* and program *j*.

### CNA estimation

Initial values (CNA_0_) were estimated by first sorting the analyzed genes by their chromosomal location and calculating a moving average of relative expression values (*ER*), with a sliding window of 100 genes, as previously described (Tirosh et al., 2016b). To avoid considerable impact of any particular gene on the moving average, in this analysis we limited relative expression values to [−3,3]. In order to define proper CNA reference values to be used as the baseline, we downloaded gene level copy number data (Affymetrix SNP6.0 arrays, log_2_(copy number/2)) from the CCLE portal (https://portals.broadinstitute.org/ccle) and calculated, for each cell line, the average copy number signal by chromosome arm. Next, for each chromosome arm, we selected a set of reference cell lines, defined as those presenting an average copy number signal ranging from −0.2 to 0.2. For a given CNA window, in a given chromosome arm, we then calculated the average CNA estimates of the respective reference cell lines and define the minimum (BaseMin) and maximum (BaseMax) values obtained as the lower and upper baseline limits. The final CNA estimate of cell *i* at position *j* was defined as:

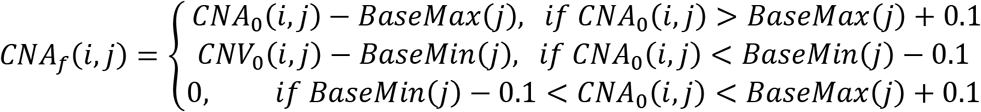

### Detection of CNA subclones within cell lines

To confidently identify CNA-based subclones we focused on CNAs encompassing whole chromosome arms, as these are more reliably inferred than focal CNAs. We reasoned that the presence of multiple subclones in a single cell line would be reflected in a multimodal distribution of CNA signal for at least one chromosome arm, across cells from a given cell line. Thus, we first calculated for each cell and each chromosome arm, the average CNA_0_ estimate across all loci in the chromosome arm. Next, we examined, for each cell line, if the distribution of arm-level CNA values was multimodal for any chromosome arm. To this end, we fitted each distribution to a Gaussian mixture model (GMM) and calculated the probability for each cell to belong to each mode by an Expectation-Maximization algorithm, implemented by the R function Mclust. A cell line was defined as having subclones if, for at least one chromosome arm, a minimum of 20 cells were classified into a second mode with at least 99% confidence. Cells were then assigned to subclones based on their mode for all chromosome arms with a multimodal distribution. CNA_0_ were used instead of CNA_f_ since the normalization procedure introduces a bias (zero inflation) in signal distribution.

### Association between programs of variability and CNA subclones

In cell lines presenting discrete programs of variability and CNA subclones, we evaluated the association between the expression-based classification of cells into subpopulations, as defined by DBSCAN, and the subclone-based classification, as defined by GMM, using Fisher’s exact test. In cell lines presenting continuous programs of variability and CNA subclones, we compared NMF cell scores of each program between clones using t-test. *P*<0.001 were considered statistically significant.

### Drug screen analysis

In order to define potential hits from the primary screen for follow-up in the secondary screen, we considered the differential viability between the EpiSen-high and EpiSen-low states for each compound in each cell line. In order to define hits that were differentially sensitive for only one cell line, we used 2.5 standard deviations from the mean of the difference in viability of the vehicle (DMSO-treated) controls between states as the threshold. In order to define shared hits that were differentially sensitive between the EpiSen-high and EpiSen-low states in both cell lines we used a less strict threshold of 2 standard deviations from the mean of the difference in viability of the vehicle controls between states. A third category of compounds that killed cells in both the EpiSen-high and EpiSen-low states at 10 µM (defined as ≤10% viability) was also selected for follow-up in the secondary screen at lower concentration (1 µM).

The secondary screen was performed in duplicates. In order to determine the differential viability between the EpiSen-high and EpiSen-low states, we compared the mean of each pair of duplicate measurements. To avoid an impact from outlier measurements, in each case where the difference between duplicates was larger than 20%, we calculated three potential values for differential viability between EpiSen-high and EpiSen-low populations: one value based on the mean of the two duplicate measurements and two additional values based on each measurement alone. We then conservatively used the minimal value of differential viability to ensure that individual outlier measurements will not lead to the appearance of differential viability. In order to define hits in the secondary screen, the threshold was defined by the upper and lower bounds of the adjusted control values over replicates between states.

A subset of compounds that were differentially sensitive between the EpiSen-high and EpiSen-low cell lines were selected for follow-up dose response studies in SCC47 in each sorted subpopulation (EpiSen-low, EpiSen-high, reference). To generate dose response curves, viability at each concentration of the seven-point dose response series was averaged over replicates and normalized to the viability of vehicle (DMSO) controls. Percent change in viability was calculated at each concentration using the normalized viability and curves were fit using these values with a three-parameter nonlinear regression model in GraphPad Prism 8 (GraphPad Software, La Jolla CA, USA) where: *Y=max%change in viability+(min%change in viability-max%change in viability)/(1+10^(X-LogEC50))* with Hill Slope=-1.0. EC50 values as determined by the three-parameter model were defined as the concentration at which the fitted curve crossed the value corresponding to half of the maximal inhibition (i.e. maximal percent change in viability). For visualization of dose response curves, normalization was performed to lowest concentration of drug. Two compounds for which EC50 could not be accurately calculated were omitted from further analysis. A paired t-test was performed using the aggregated differences in viability at each concentration to determine statistical significance (*P*≤0.05) of differential viability between curves.

## Data availability

Raw and processed scRNA-seq data will be available through the Gene Expression Omnibus (GEO) and the Broad Institute’s single cell portal.

## Acknowledgements

This work was supported by funding from the Human Frontiers Science Program (I.T.), the Israel Science Foundation (I.T.), the Zuckerman STEM leadership program (I.T.), the Mexican Friends New Generation grant (I.T.), the Rising Tide Foundation (I.T.), the A.M.N. Fund for the Promotion of Science, Culture and Arts in Israel (I.T.), the Estate of Dr. David Levinson, the Dr. Celia Zwillenberg-Fridman and Dr. Lutz Zwillenberg Career Development Chair (I.T.), the Sao Paulo Research Foundation (FAPESP) Fellowship 2014/27287-0 and 2017/24287-8 (G.S.K.), the Clore Foundation Postdoctoral Fellowship (A.C.G.), the Klarman Cell Observatory (A.R.) and HHMI (A.R.).

## Competing interests

A.R. is a founder and equity holder of Celsius Therapeutics and a SAB member of Neogene Therapeutics, ThermoFisher Scientific and Syros Pharmaceuticals.

