## Supplemental Figures for "Pan-cancer single cell RNA-seq uncovers recurring programs of cellular heterogeneity"

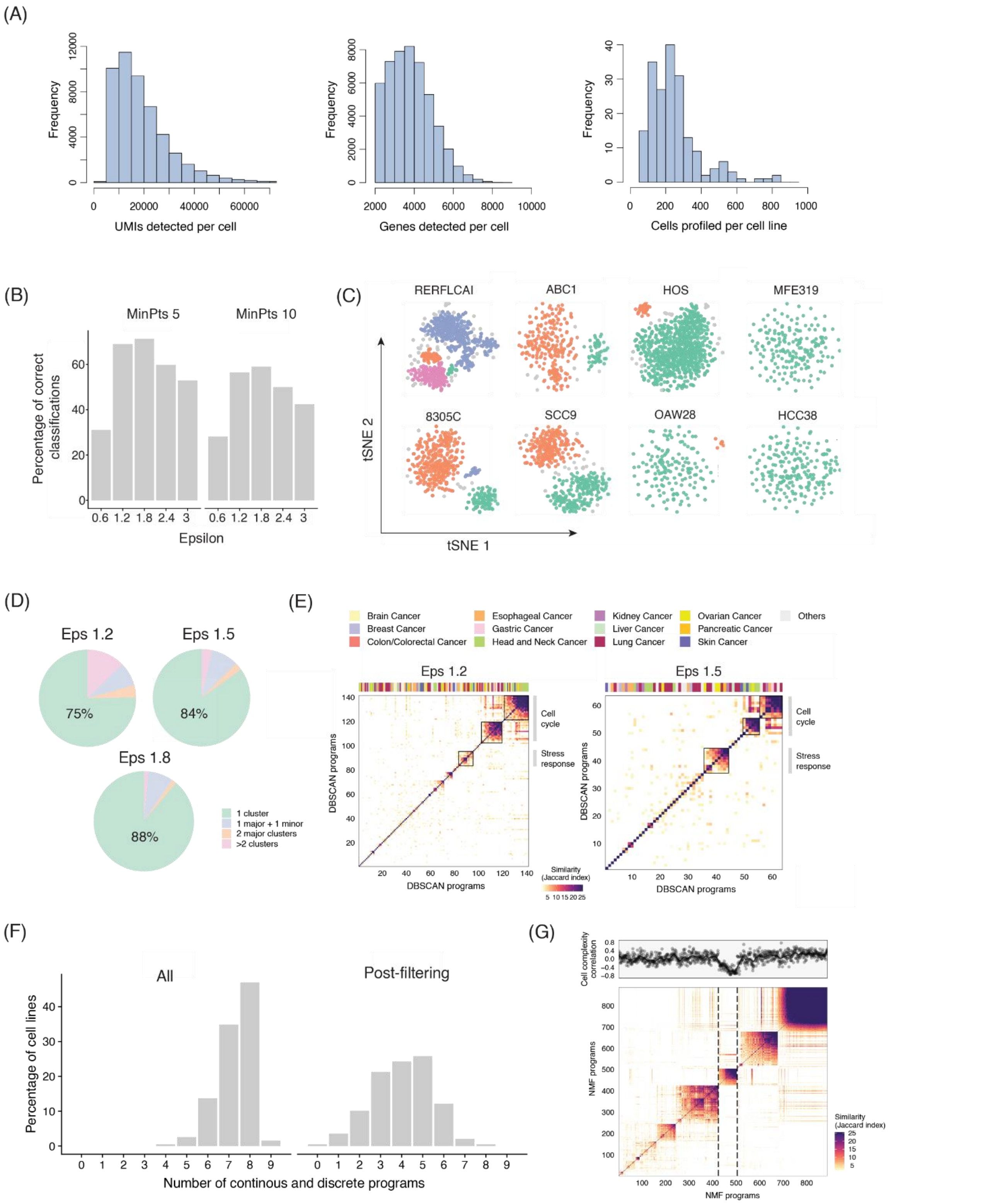

**Figure S1. Quality controls and analysis of discrete and continuous heterogeneity**. (**A)** Quality controls for multiplexed scRNA-seq of cell lines. Histograms show distributions of the number of UMIs detected per cell (left), the number of genes detected per cell (middle), and the number of cells detected per cell line (right). **(B)** Performance of DBSCAN using different sizes of epsilon neighborhood (eps) and minimum numbers of points required to form a dense region (MinPts). We randomly selected cells from two different cell lines and tested the ability of DBSCAN to distinguish between them using different parameter combinations. The procedure was repeated 1,000 times and the combination yielding the highest rate of correct classification was applied in the subsequent analyses. (**C)** t-SNE plots for additional two examples of cell lines from each of the four classes defined by presence and number of discrete subpopulations identified by DBSCAN (as in **Fig. 2B**). **(D-E)** Identification of discrete programs of heterogeneity, as in **Fig. 2B**, using less stringent eps (1.2 and 1.5) highlights common trends. **(F)** Number of heterogeneity programs identified per cell line using NMF. NMF was applied to each cell line using k (number of factors) of 6-9, and gene programs identified as variable with 2 or more values of k were retained (left panel). To identify common expression programs varying within multiple cell lines, we excluded programs with limited similarity to all other programs as well as those associated with technical confounders (right panel). (**G)** Pairwise similarities between programs identified by NMF across all the cell lines analyzed, with cell lines ordered by hierarchical clustering. Programs with limited similarity to all other programs were excluded. Top panel indicates correlations between program scores and cell complexity (i.e. number of genes detected per cell). The cluster of programs that correlates with complexity (indicated by dashed lines) was excluded from subsequent analyses.

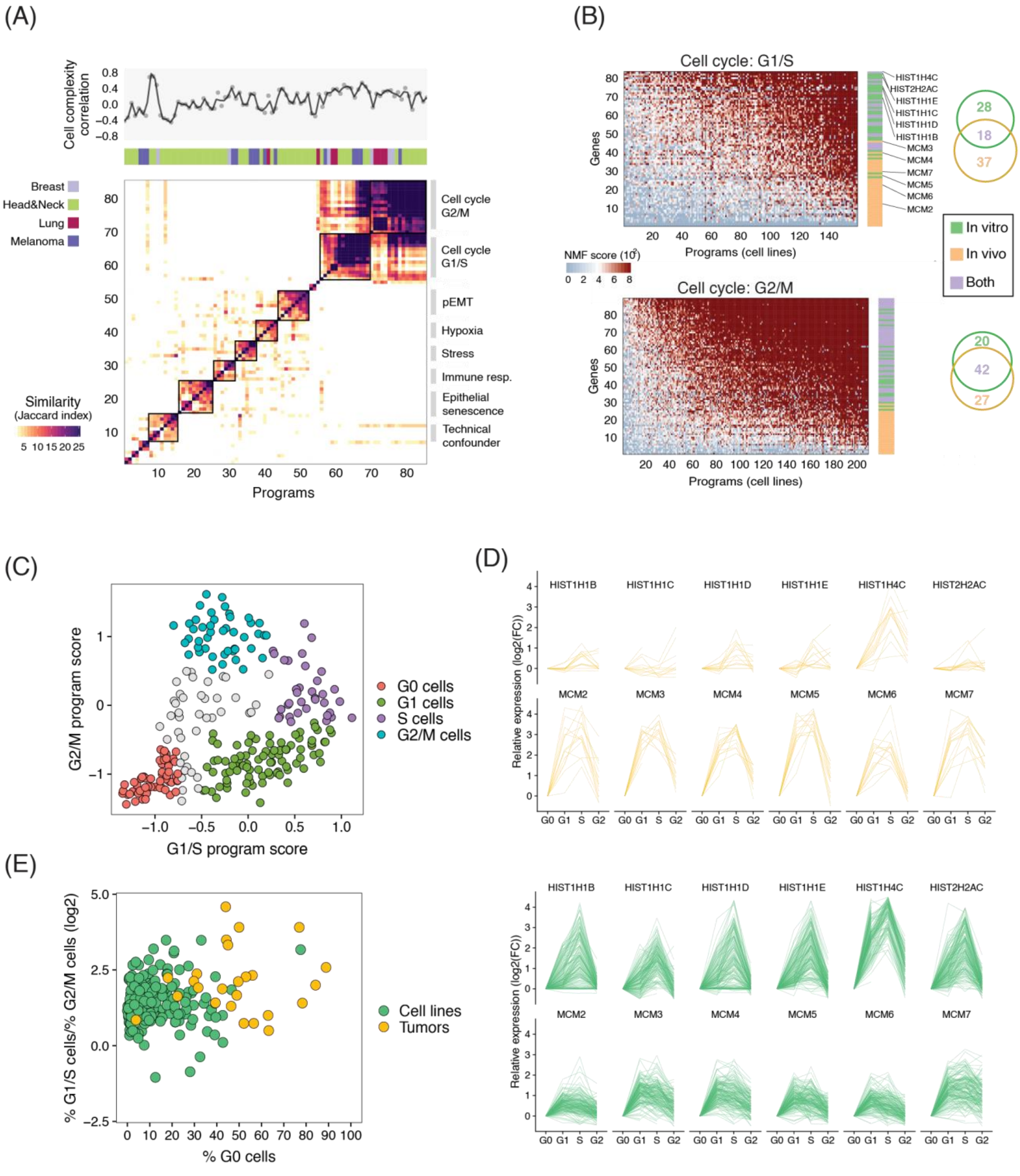


**Figure S2. Comparison between cell cycle in vitro and in vivo.** (**A**) Heatmap depicts pairwise similarities between programs identified in tumor samples using NMF. Programs with limited similarity to all other programs were excluded. Top panel shows tumor type and correlations between program scores and cell complexity (i.e. number of genes detected per cell). Hierarchical clustering emphasizes multiple clusters (shown by squares), one of which is correlated with cell complexity and thus excluded as a potential technical artifact. (**B**) NMF scores of G1/S genes (top panel) and G2/M genes (bottom panel) across all NMF programs associated with the corresponding cell cycle phase; each program is from a different cell lines. Genes are ranked in each panel by average scores, and their assignment to *in vitro* and *in vivo* cell cycle programs is indicated in the right bar, demonstrating that G1/S programs differ both across cell lines and between cell lines and tumors, while G2/M programs are more consistent. Venn diagrams (right) illustrate the overlap of genes between *in vivo* and *in vitro* RHPs. (**C**) Single-cell profiles showing G1/S and G2/M program score thresholds used to assign cells to different cell cycle phases. (**D**) Examples of genes with distinct cell cycle upregulation *in vitro* and *in vivo*. Expression of HIST genes (preferentially induced *in vitro*) and MCM genes (preferentially induced *in vivo*) is shown along the cell cycle (relative to cells in G0) in cell lines (C, green lines) and tumors (D, yellow lines). (**E**) Comparison of cell cycle phase distribution *in vitro* and *in vivo*. Scatterplot shows the percentage of cells in G0 (x-axis) and the ratio between the percentage of cells in G1/S and G2/M (y-axis) for each cell line (green) and tumor (yellow) analyzed. Cell lines display a significantly lower percentage of cells in G0 cells ($p=2e^{-10}$, t-test).

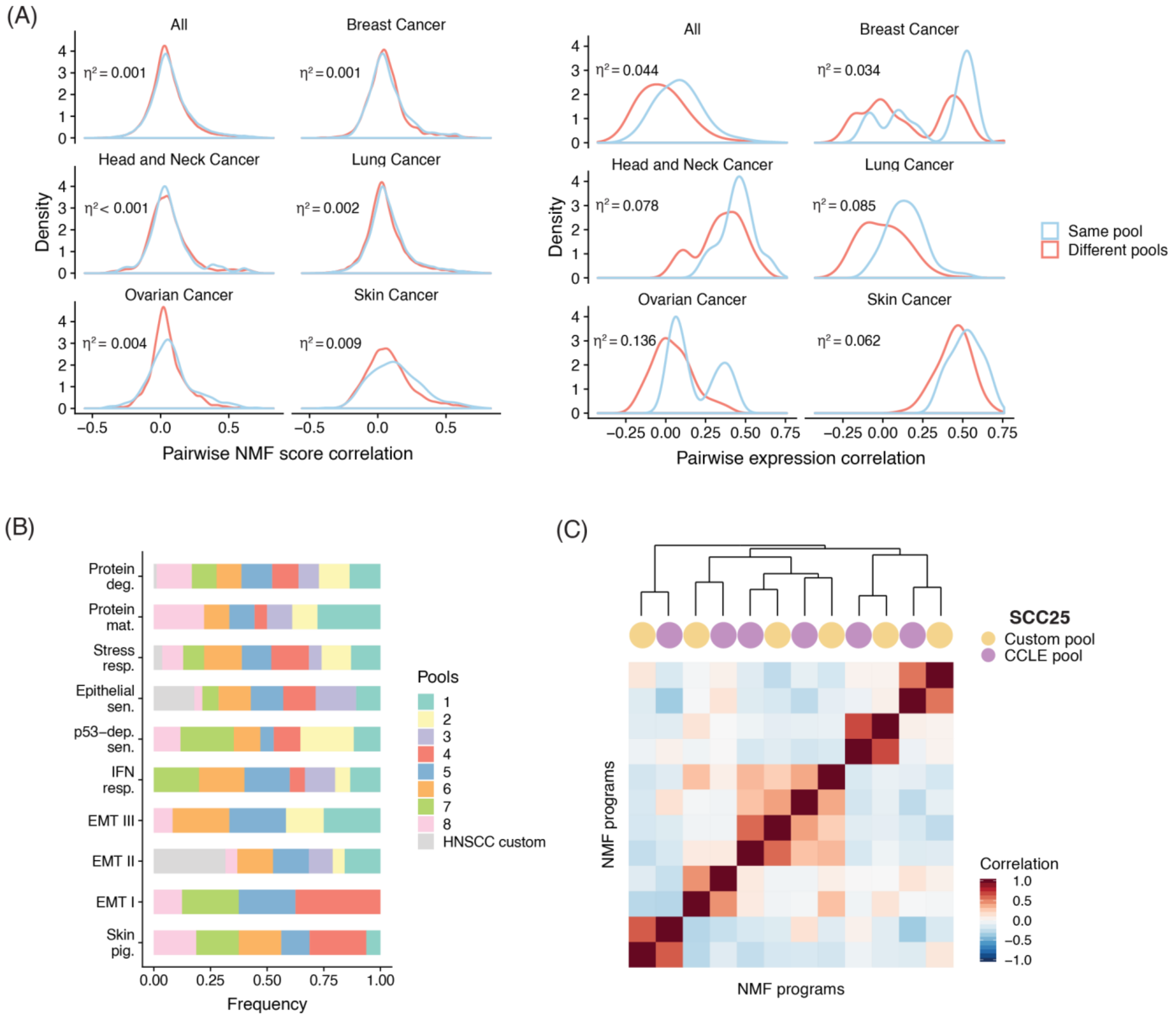


**Figure S3. The pooling procedure has a limited impact on expression heterogeneity within cell lines. (A)** Distribution of pairwise similarities between expression patterns of cell lines in the same (blue) and in different (red) pools. Left: The comparison was done for all NMF programs detected in the corresponding cell lines, by correlation of the NMF gene scores (for all analyzed genes). Right: The comparison was done between the average expression profiles of the cell lines, by correlation across all analyzed genes. Comparisons were performed across all cell lines or separated by cancer type (only most abundant types are shown). The proportion of total variance ($\eta^2$) explained by whether or not programs/cell lines were in the same pool (calculated using one-way ANOVA) is indicated, suggesting that patterns of expression heterogeneity (left) are largely unaffected by the pool microenvironment, while the average expression profiles (right) are more affected. **(B)** Distribution of the pool of origin of RHPs. Each RHP was observed in multiple pools, underscoring the lack of pool-specific effects. **(C)** Pairwise correlations between NMF programs obtained for the HNSCC cell line SCC25, which is the only cell line that was profiled in two different pools. Hierarchical clustering reveals highly concordant programs in the two pools.

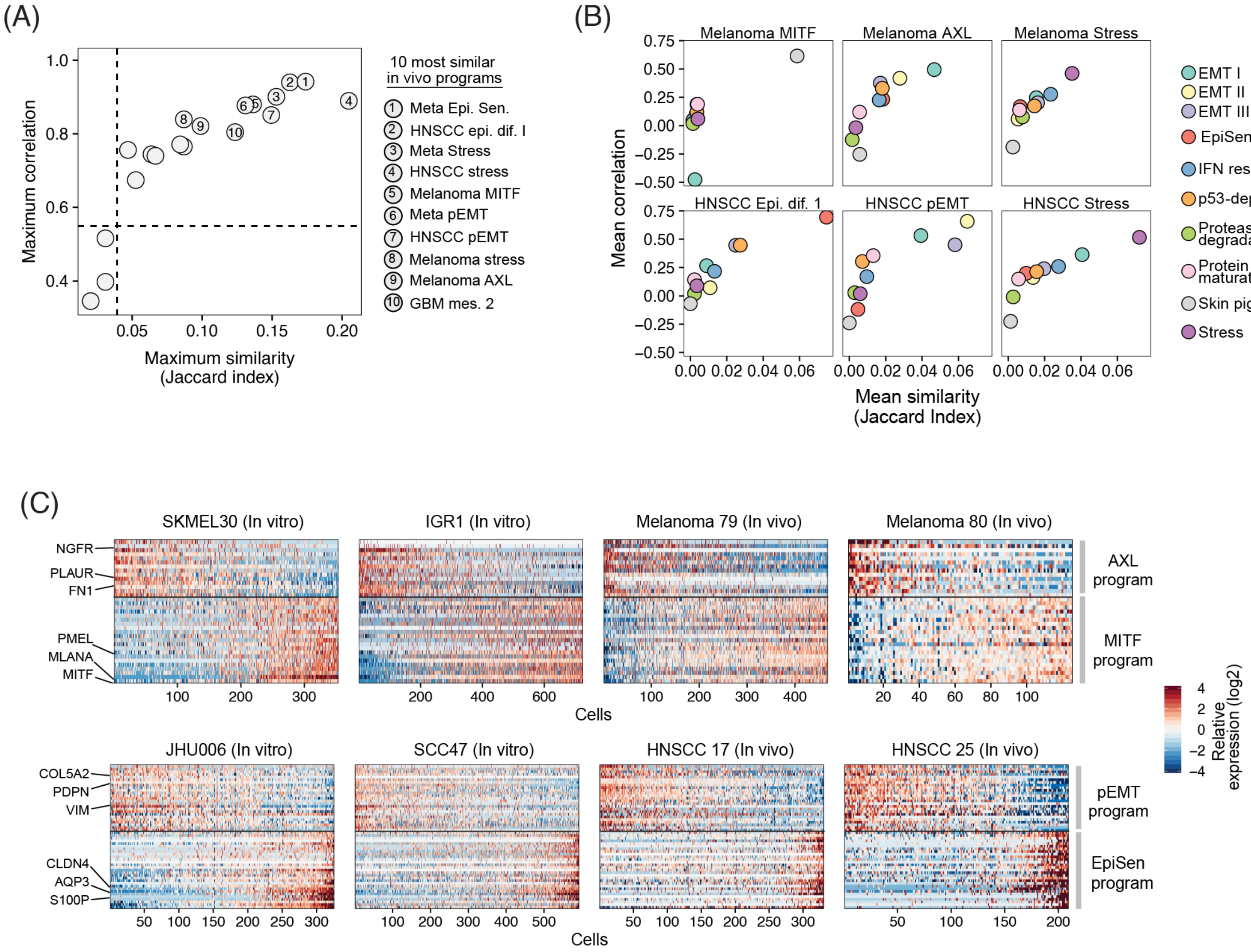


**Figure S4. Cell lines recapitulate programs of heterogeneity observed in tumor samples. (A)** Scatterplot shows maximum correlation of single-cell score (Y-axis) and gene overlap (X-axis) between each *in vivo* program and the *in vitro* programs shown in **Fig. 3A**. The top 10 in vivo programs with highest correlations are annotated, and relevant ones are also highlighted in **Fig. 3B top**. **(B)** Scatterplots show mean single-cell score correlations (Y-axis) and mean similarities (X-axis) between relevant in vivo programs and the 10 in vitro RHPs. **(C)** Heatmap shows relative expression of genes shared by paired *in vivo* and *in vitro* programs in selected melanoma and HNSCC cell lines and tumors, highlighting similar patterns of variability *in vivo* and *in vitro*. Cells are sorted according to the relative average expression of genes in each program, showing the negative correlation between the AXL and MITF programs in melanomas and the pEMT and EpiSen programs in HNSCC. Programs are annotated (right) and selected genes are indicated (left).

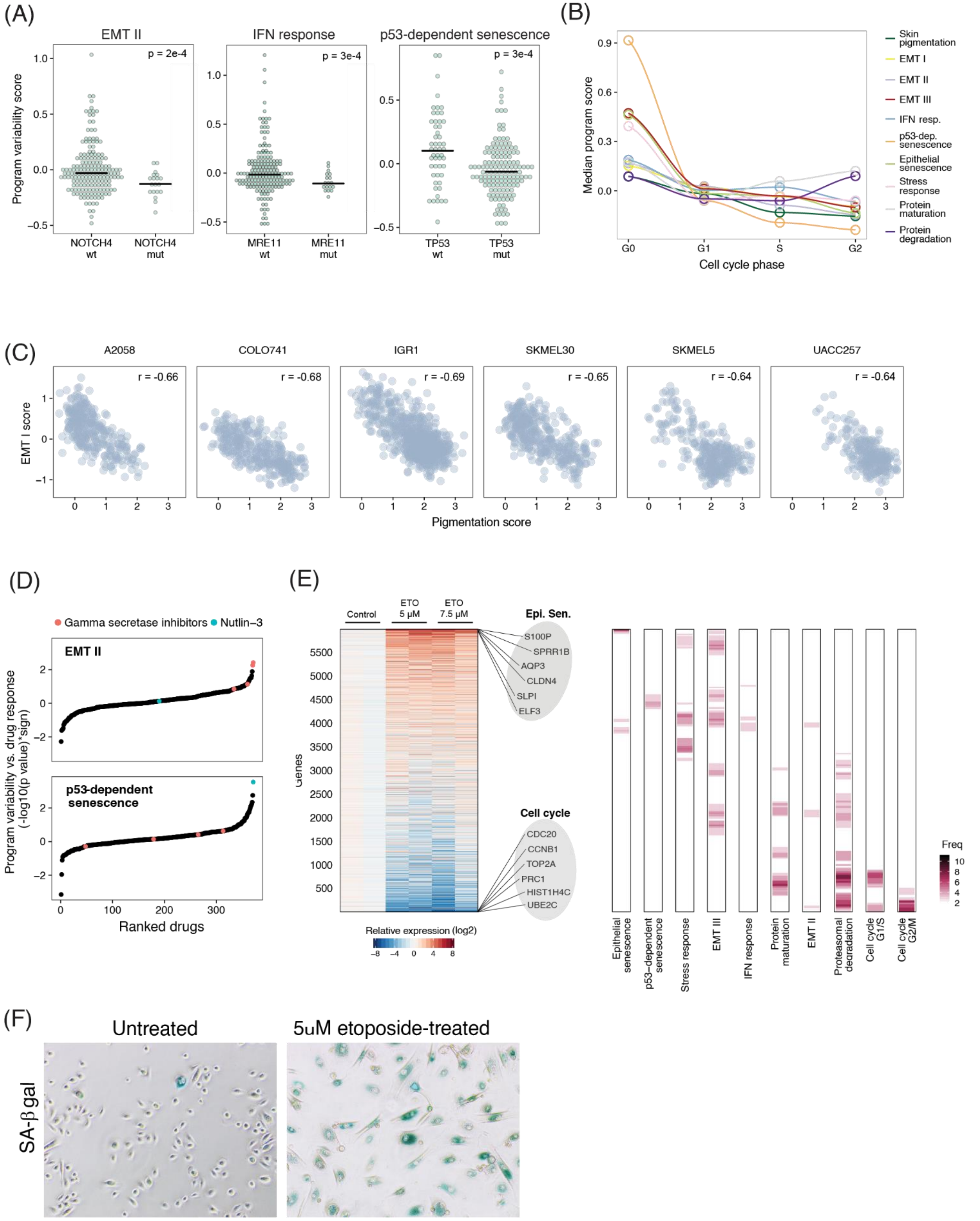
(A)
EMT II
IFN response
p53-dependent senescence
p = 2e-4
p = 3e-4
p = 3e-4
Program variability score
NOTCH4 wt
NOTCH4 mut
MRE11 wt
MRE11 mut
TP53 wt
TP53 mut
(B)
Median program score
Cell cycle phase
G0
G1
S
G2
Skin pigmentation
EMT I
EMT II
EMT III
IFN resp.
p53-dep. senescence
Epithelial senescence
Stress response
Protein maturation
Protein degradation
(C)
A2058
COLO741
IGR1
SKMEL30
SKMEL5
UACC257
r = -0.66
r = -0.68
r = -0.69
r = -0.65
r = -0.64
r = -0.64
EMT I score
Pigmentation score
(D)
Gamma secretase inhibitors
Nutlin–3
EMT II
p53-dependent senescence
Program variability vs. drug response (-log10(p value)*sign)
Ranked drugs
(E)
Control
ETO 5 μM
ETO 7.5 μM
Epi. Sen.
S100P
SPRR1B
AQP3
CLDN4
SLPI
ELF3
Cell cycle
CDC20
CCNB1
TOP2A
PRC1
HIST1H4C
UBE2C
Genes
Relative expression (log2)
Freq
Epithelial senescence
p53–dependent senescence
Stress response
EMT III
IFN response
Protein maturation
EMT II
Proteasomal degradation
Cell cycle G1/S
Cell cycle G2/M
(F)
Untreated
5uM etoposide-treated
SA-β gal

**Figure S5. Determinants and consequences of cellular heterogeneity.** (**A**) Association between RHP variability scores and somatic non-silent mutations. We compared the variability of each program in mutated and non-mutated cell lines using t-test. Model cell lines (high variability) of EMT-II, IFN response and p53-dependent senescence RHPs are depleted of NOTCH4, MRE11 and TP53 mutations, respectively. (**B**) Median RHP scores of cells in each phase of the cell cycle, emphasizing the high expression of the senescence, stress and EMT-III metaprograms in non-cycling cells (G0). Cell cycle state was estimated for each individual cell based on the relative expression of the G1/S and G2/M metaprograms. For each RHP we only considered the respective model cell lines. (**C**) Single-cell profiles show a negative correlation between the skin pigmentation and the EMT-I RHPs within six melanoma cell lines. (**D**) Association between drug response (CTRP database) and program variability calculated using linear regression including tumor type and program variability as independent variables. Increased sensitivity to NOTCH inhibition (gamma secretase inhibitors) and MDM2 inhibition (Nutlin-3) were observed in model cell lines (high variability) of the EMT-II and the p53-dependent senescence respectively. (**E**) Heatmap depicts the relative expression of 6,000 genes (rows) in primary lung bronchial cells, 9 days after induction of senescence by etoposide treatment for 48h in two concentrations. Bars on the right show the frequency of RHP signature genes within sliding windows of 300 genes. RHPs are sorted from left to right by their enrichment with upregulated and downregulated genes, respectively. EpiSen and cell cycle programs were the two extreme programs, and selected genes from these programs are labeled. (**F**) Induction of senescence in primary lung bronchial cells confirmed by SA-β-gal staining.

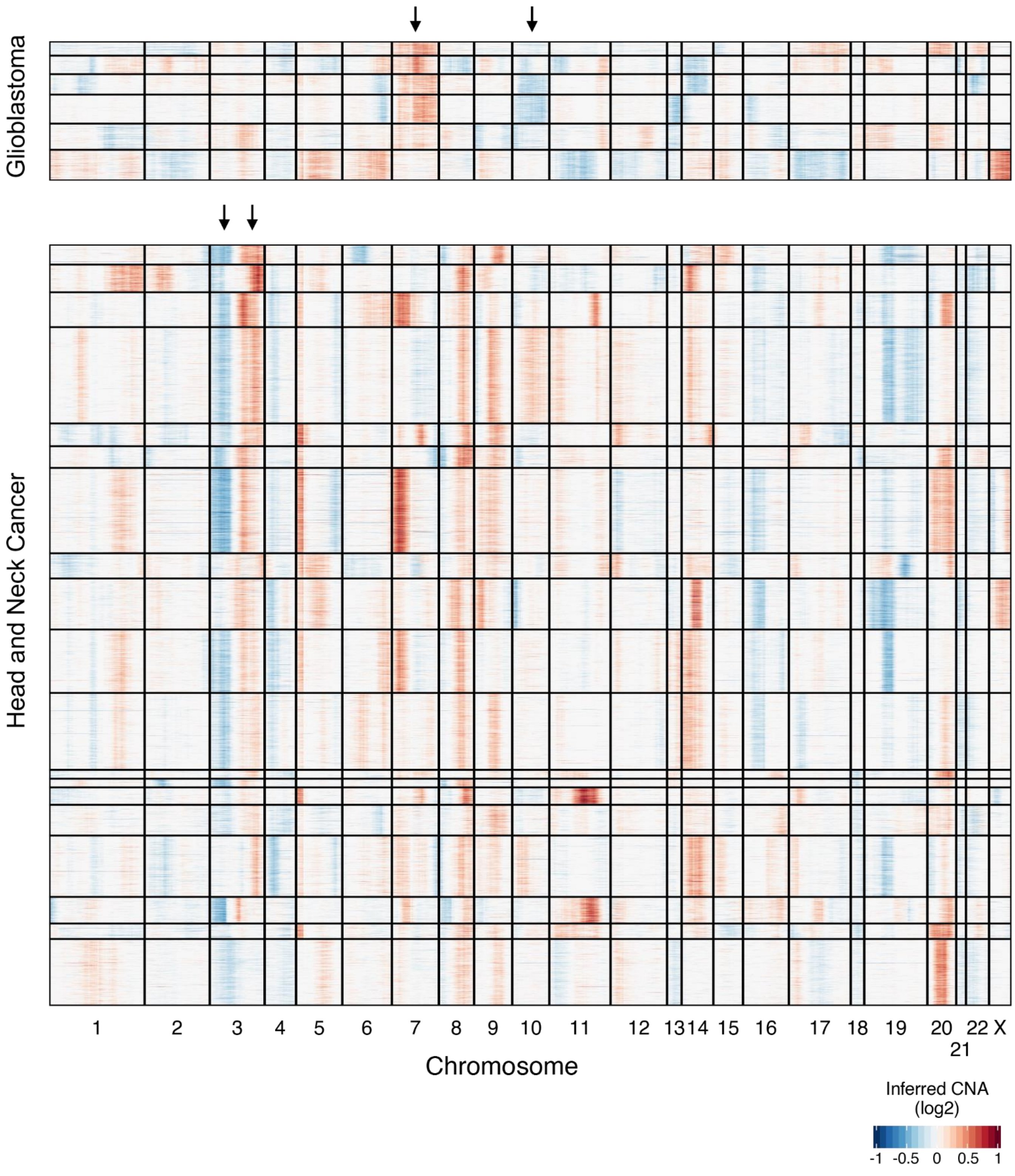


**Figure S6. Inferred CNAs are consistent with expected chromosomal aberrations**. Heatmap depicts inferred CNAs for individual cells (rows) from 6 glioblastoma (top) and 19 HNSCC (bottom) cell lines, based on average expression in sliding windows of 100 genes. Arrows highlight expected hallmark alterations - the gain of chromosome 7 and loss of chromosome 10 in glioblastoma, and the loss of chromosome 3p and gain of chromosome 3q in HNSCCs.

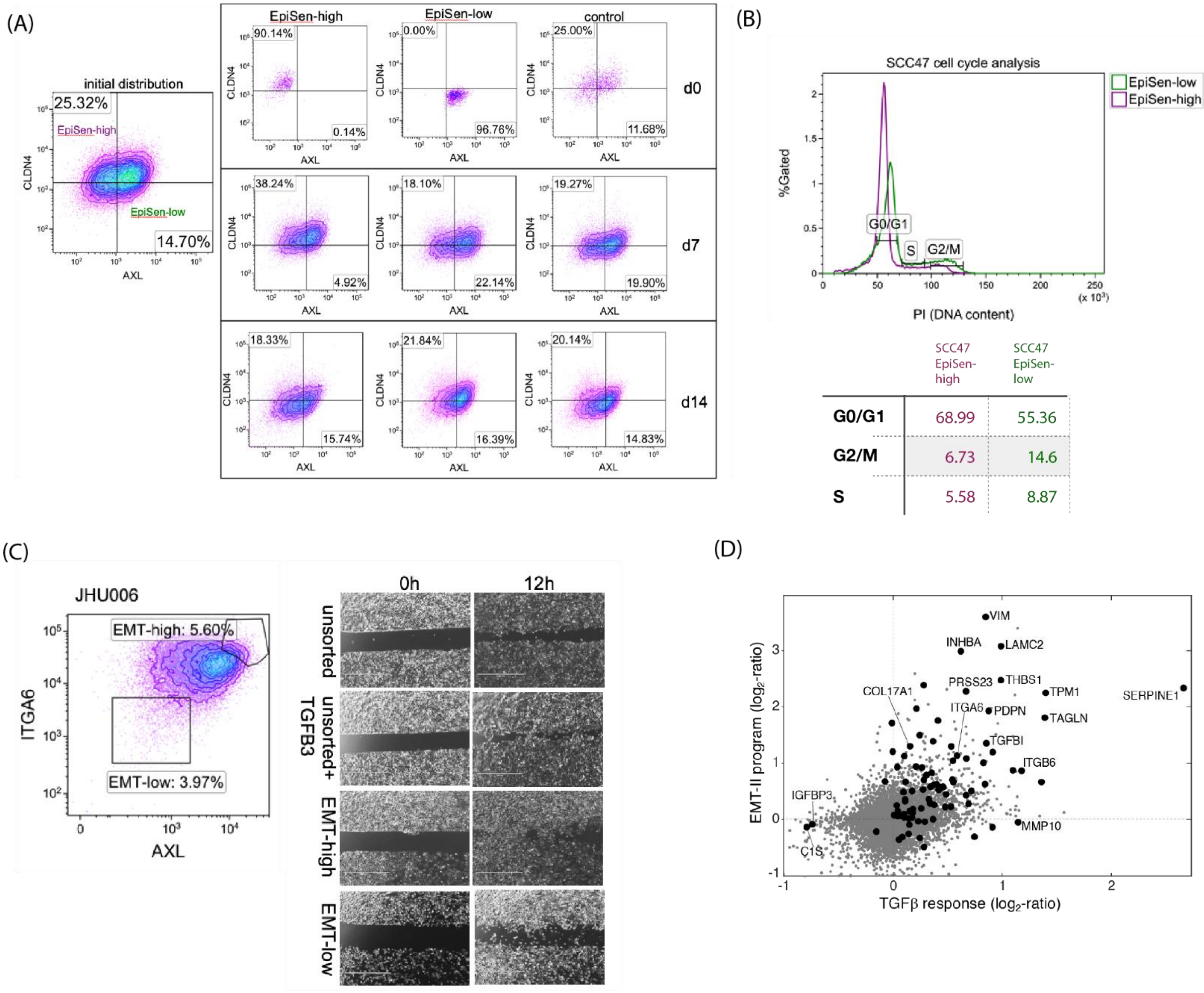


**Figure S7. Interrogating EpiSen and EMT-II in model cell lines.**
(**A**) Three JHU006 subpopulations were isolation by FACS (EpiSen-high: AXL$^-$CLDN4$^+$, EpiSen-low population: AXL$^+$CLDN4$^-$, control: unsorted), and analyzed immediately after sorting (day 0, top) and at two additional time points in culture (day 7 and 14, middle and bottom, respectively). Density plots correspond to the pie charts in Fig. 5C. (**B**) FACS analysis of cell cycle by the DNA binding dye propidium iodide (PI) on sorted EpiSen-high and EpiSen-low cells in SCC47, as shown for JHU006 in Fig. 5D. The table below summarizes the results. (**C**) Left: isolation by FACS of the EMT-II-high population (AXL$^+$ITGA6$^+$) and the EMT-II-low population (AXL$^-$ITGA6$^-$) in JHU006. Right: gap closure (migration) assay on unsorted, unsorted but TGFβ–3-treated, EMT-II-high, and EMT-II-low cells at 0h and 12h following gap generation. **(D)** Comparison of EMT program induced upon TGFβ treatment of unsorted cells (X-axis) vs. the EMT-II RHP gene scores (Y-axis). In both axes, data was averaged over the results for JHU006 and SCC47.

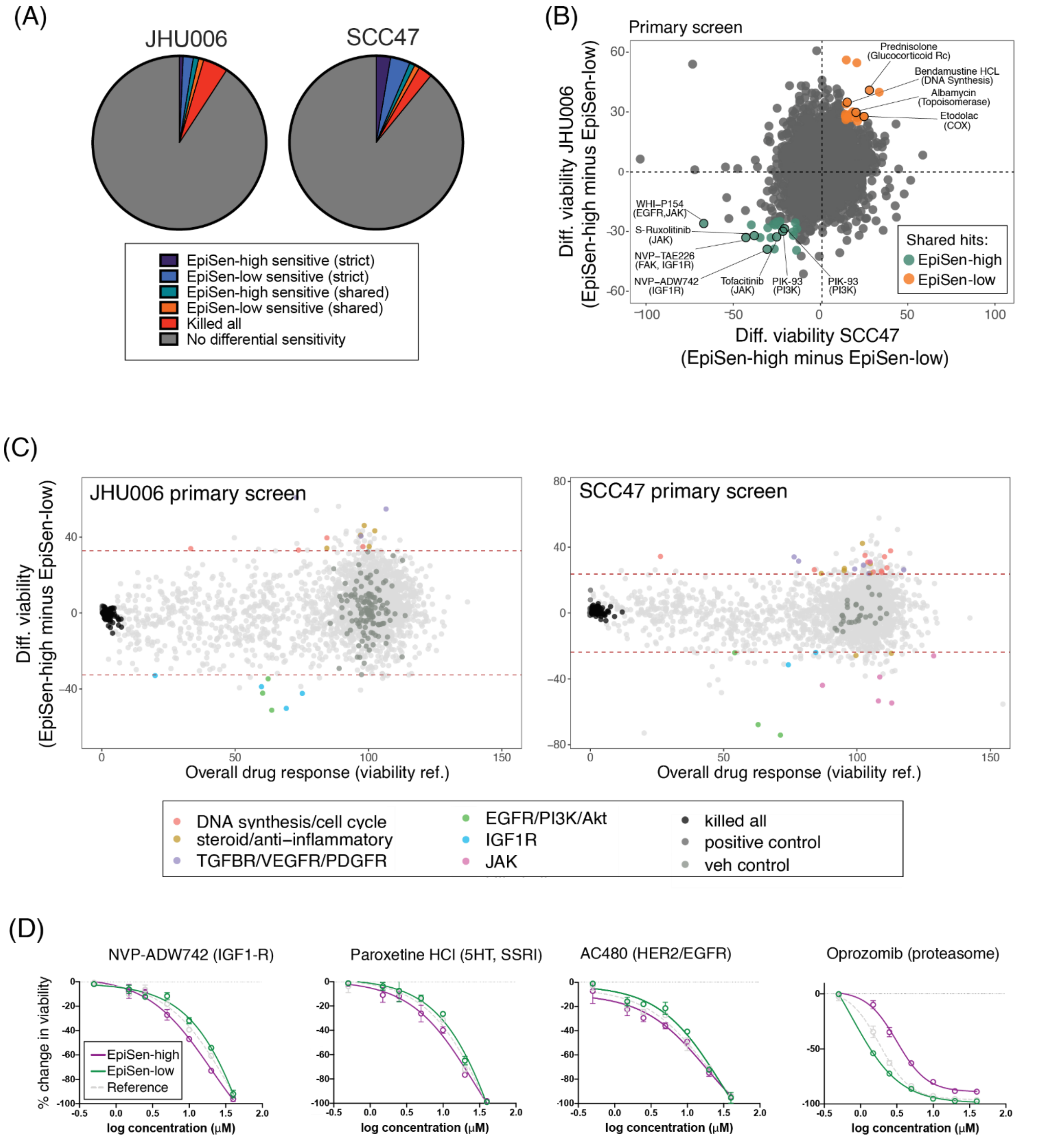


**Figure S8. Drug sensitivity of EpiSen-high cells and EpiSen-low cells**. (**A**) Pie charts depict the proportions of primary screen hits by type. (**B**) Shared hits between SCC47 and JHU006 for compounds that preferentially killed the EpiSen-high (green) and EpiSen-low (orange) states. Selected hits are labeled (**C**) Viability of the control population (X-axis) and differential viability of the EpiSen-high vs. EpiSen-low populations (Y-axis) upon treatment with 2198 compounds in JHU006 (left) and SCC47 (right). Dotted lines represent thresholds for differential sensitivity, and hits are colored as defined in the lower legend. (**D**) Dose response curves of selected compounds in three SCC47 subpopulation (continued from Fig. 6C). Change in viability was calculated relative to vehicle (DMSO-treated) controls. Error bars represent standard deviation.
